# Unveiling unexpected complexity and multipotentiality of early heart fields

**DOI:** 10.1101/2021.01.08.425950

**Authors:** Qingquan Zhang, Daniel Carlin, Fugui Zhu, Paola Cattaneo, Trey Ideker, Sylvia M. Evans, Joshua Bloomekatz, Neil C. Chi

## Abstract

Complex organs are composed of a multitude of specialized cell types which assemble to form functional biological structures. How these cell types are created and organized remains to be elucidated for many organs including the heart, the first organ to form during embryogenesis. Here, we show the ontogeny of mammalian mesoderm at high-resolution single cell and genetic lineage/clonal analyses, which revealed an unexpected complexity of the contribution and multi-potentiality of mesodermal progenitors to cardiac lineages creating distinct cell types forming specific regions of the heart. Single-cell transcriptomics of *Mesp1* lineage-traced cells during embryogenesis and corresponding trajectory analyses uncovered unanticipated developmental relationships between these progenitors and lineages including two mesodermal progenitor sources contributing to the first heart field (FHF), an intraembryonic and a previously uncharacterized extraembryonic-related source, that produce distinct cardiac lineages creating the left ventricle. Lineage-tracing studies revealed that these extraembryonic-related FHF progenitors reside at the extraembryonic-intraembryonic interface in gastrulating embryos and generate cardiac cell types that form the epicardium and the dorsolateral regions of the left ventricle and atrioventricular canal myocardium. Clonal analyses further showed that these progenitors are multi-potent, creating not only cardiomyocytes and epicardial cell types but also extraembryonic mesoderm. Overall, these results reveal unsuspected multiregional origins of the heart fields, and provide new insights into the relationship between intraembryonic cardiac lineages and extraembryonic tissues and the associations between congenital heart disease and placental insufficiency anomalies.

## Introduction

Embryonic development is a process by which a single cell with potential to give rise to all cells within the embryo progressively creates groups of cells with more restricted potential. A developmental field is a collection of cells with a shared potential to produce a restricted subset of embryonic structures. By their nature, developmental fields are transient and present only at specific developmental stages. Studies of heart development over the last few decades have defined a first heart field (FHF) and a second heart field (SHF), according to their potential to give rise to specific myocardial lineages within the developing heart^1^.

The FHF and SHF were inferred by retrospective clonal analyses in the mouse embryo, which revealed two clonally distinct differentiated myocardial lineages, the first and second heart lineages, respectively^2–4^. At E8.5, clones of the first heart lineage were observed to be excluded from the outflow tract, populate the entire left ventricle (LV) and left atrioventricular canal (AVC), and contribute some cells to both atria and right ventricle (RV), whereas clones of the second heart lineage were found to be excluded from the LV^2^. Of the heart fields predicted by this model, the SHF has been visualized and defined, as a population of cells medial to the differentiating myocardial cells of the cardiac crescent that expresses the transcription factor Isl1 around E7.75^5^. SHF cells expressing Isl1 will also give rise to multiple other cell types that contribute to the heart, pharyngeal arches and head/neck including endothelial, endocardial and smooth/skeletal muscle cells^5–11^. At E7.75, the first differentiating cells in the cardiac crescent are marked by the ion channel *Hcn4*. As *Hcn4-CreERT2* labeled cells in the crescent mainly contribute to cardiomyocyte lineages in the LV and parts of the atria, they are thought to represent more differentiated precursors of first heart lineage cardiomyocytes, and for that reason have been considered as representatives of the FHF at crescent stages^12,13^. However, the origins and attributes of FHF progenitors prior to cardiac crescent stages remain unknown.

In addition to myocardial and endocardial lineages, the fully formed heart includes fibroblasts and vascular support cells that derive from the epicardium. The proepicardium, a transient cluster of cells that forms at the base of the looping heart from the septum transversum (ST) during early heart development, produces cells which cover the heart surface as an epithelium to form the epicardium^14^. Subsets of cells from the epicardium will migrate into the myocardium to give rise to cardiac fibroblasts and vascular support cells of the coronary vasculature, which are essential for formation and function of the heart^15–17^. Yet, the developmental origin of the proepicardium, and its relationship to previously described heart fields remains to be defined^18^.

To address the developmental origins, definition and contribution of specific cell lineages creating the heart, we performed single cell transcriptomic analyses on *Mesp1-cre; Rosa26-tdTomato (Rosa26-tdT)* mouse embryos across key developmental stages of cardiac development. Computational trajectory analyses of these data notably predicted a potential group of progenitors specifically expressing *Hand1* that may give rise to a subset of first heart lineage cardiomyocytes. Notably, *in situ* and lineage tracing analyses utilizing *Hand1-CreERT2* revealed a *Hand1*-expressing population at the extraembryonic/intraembryonic boundary of the gastrulating embryo that contributes to first heart lineage cardiomyocytes residing largely within dorsolateral regions of the LV and AVC. Intriguingly, the *Hand1-CreERT2* lineage created only a subset of (rather than all) LV cardiomyocytes. As the second heart lineage does not populate the LV^2^, this finding implies a previously unexpected complexity of the FHF in which the FHF is not a single developmental heart field, but rather composed of at least two distinct developmental heart fields, one of which, identified here, is marked by *Hand1*. Earlier studies have presumed that the FHF, in contrast to the SHF, has a tightly restricted developmental potential, only giving rise to myocardial cells within specific segments of the heart. Utilizing *Hand1-CreERT2* and *Rosa26-Confetti* clonal analyses, we surprisingly found that the *Hand1-CreERT2*-marked FHF is composed of multipotent cells that give rise to not only first heart lineage myocardial cells, but also serosal mesothelial lineages (including proepicardium/epicardium and pericardium), and cells within extraembryonic mesoderm.

Overall, our results reveal a closer lineage relationship than previously suspected between cardiac tissues and extraembryonic mesoderm. Our observation that the *Hand1* expressing segment of the FHF gives rise to epicardial cells also provides insight into the developmental origins of the epicardium, thus uncovering a new early clonal relationship between cardiac muscle cells of the first heart lineage and cells of the epicardium.

## Results

### scRNA-seq analysis of *Mesp1* lineage-traced cells reveals developmental cell types participating in mesoderm-related organogenesis

As *Mesp1* is known to mark early mesoderm, we employed a mouse *Mesp1-*Cre^19^; *Rosa26-*tdTomato (*R26R-tdT*)^20^ genetic fate mapping system to permanently label and track all cell lineages contributing to the development of mesoderm-derived organs including the heart ^19,21–24^ (Fig. 1**a**). To discover the broad spectrum of developmental cell types participating in this process, we interrogated the transcriptomes of individual *Mesp1-Cre; Rosa26-tdT* genetically-labeled cells utilizing single-cell RNA-sequencing (scRNA-seq) (Fig. 1**a**). Because of our focus on early mesoderm-related organogenesis, we specifically examined isolated *Mesp1-Cre; Rosa26-tdT* single cells at E7.25 (no bud stage), E7.5 (early bud stage), E7.75 (late head fold stage) and E8.25 (somite stage) (Fig. 1**a**, Extended Data Fig. 1). Each sample was processed and analyzed through our standard pipeline and confirmed for replicate reproducibility (Extended Data Fig. 1**a-c**). t-distributed stochastic neighbor embedding (tSNE) visualization and unsupervised k-means clustering^25^ of these combined single cell data revealed a broad array of cell types, which were identified based on gene expression, during mesoderm development (Fig. 1**b-f**, Extended Data Fig. 1 and 2, Supplementary Table. 1). As mesodermal progenitors differentiated into organ-specific cell types during mouse embryogenesis, we observed that the number of identified cell-types increased with developmental age. For example, nascent, early-extraembryonic and hemogenic mesoderm (NM, EEM, Hem) cell types were detected at E7.25 as previously described^26,27^; however, many more intermediate and differentiated organ-specific cell types were identified by E7.75 and E8.25, including two cardiomyocyte clusters – developing cardiomyocytes (DC) and cardiomyocytes (CM) (Fig. 1**b**). These cell clusters appear to represent early (developing) and more established cardiomyocytes, respectively, based on their differential expression of sarcomeric (*Tnnt2*, *Ttn* and *Myl3*) and cardiac progenitor genes (*Tbx5, Sfrp1/5* and *Meis1*) (Fig. 1**f**).

**Figure 1.**
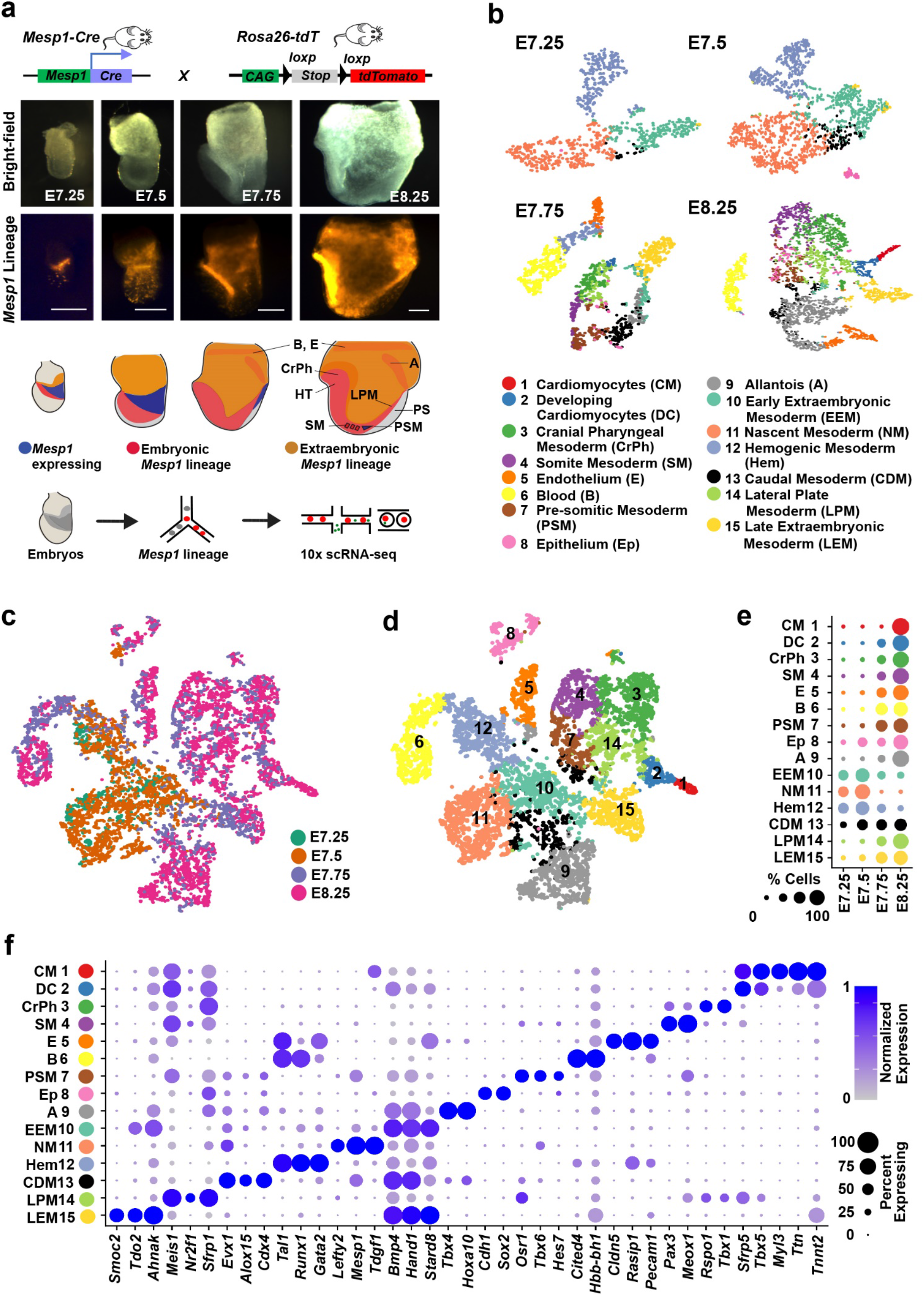
*Mesp1*-Cre single-cell maps reveal diverse cell types participating in early mouse mesoderm development. **a**, *Mesp1*-Cre scRNA-seq experimental design. *Mesp1-Cre*; *Rosa26-tdT* embryos were harvested for scRNA-seq at E7.25 (no bud stage); E7.5 (early bud stage); E7.75 (early head fold stage); and E8.25 (somite stage) as shown in representative bright-field and *Mesp1*-Cre; tdT+ (*Mesp1* lineage) micrographs. Illustration below these micrographs shows tissues genetically labeled by *Mesp1-Cre* in embryos, and workflow for capturing these labeled single cells for RNA sequencing. Scale bars, 150 μm. **b**, scRNA-seq data is displayed by tSNE plots at each developmental stage. Cells are colored according to their cell identities in **d**, **e**, **f**. **c**, **d**, tSNE plot of scRNA-seq data across all examined stages displays individual cells (single dots) by (**c**) developmental stages or (**d**) cell types. **e**, Dot plot shows distribution of each cell type across different embryonic stages. **f**, Dot plot of key marker genes identifies each cell cluster. A, Allantois; B, Blood; CDM, Caudal Mesoderm; CrPh, Cranial-pharyngeal mesoderm; CM, Cardiomyocytes; DC, Developing Cardiomyocytes; E, Endothelium; Ep, Epithelium; EEM, Early Extraembryonic Mesoderm; Hem, Hemogenic Mesoderm; HT, Heart tube; LEM, Late Extraembryonic Mesoderm; LPM, Lateral plate mesoderm; NM, Nascent Mesoderm; PSM, Pre-somitic mesoderm; PS, Primitive streak, SM, Somite mesoderm.

### Trajectory analysis elucidates developmental pathways during mesoderm organogenesis

To illuminate the developmental origins and cell fate decisions of organ-specific cell types arising from mesodermal progenitor cells including cardiac cell types, we organized cells from our single cell data along developmental trajectories using the lineage inference analysis, URD^28^, which is based on a random walk of the nearest neighbor graph of gene expression. These reconstructed developmental trajectories, as displayed in the tree structure that URD produces, not only ordered cells by a pseudotime which correlate with the developmental age of analyzed cells but also revealed both new and known developmental cell fate decisions (Fig. 2**a**, Extended Data Fig. 3**a**). In particular, we observed developmental trajectories that identified previously described mechanisms of development for some discovered cell types including the early differentiation and bifurcation of endothelial and blood cells^29^, the differentiation of somitic mesoderm from a pre-somitic state originating in the caudal mesoderm^30,31^ and a common progenitor that gives rise to cranial pharyngeal, lateral plate mesoderm and cardiomyocytes^1,9,11,32^ (Fig. 2**a**). On the other hand, examination of the cardiomyocyte developmental trajectory uncovered two potential developmental sources that may contribute to developing cardiomyocytes: a known intraembryonic progenitor from the lateral plate mesoderm (LPM) and a previously undescribed cardiac progenitor from the late extraembryonic mesoderm (LEM) (Fig. 2**a**, box). Viewing these developmental trajectories as a three-dimensional force-directed URD representation revealed how these two progenitor sources originate and then converge to independently contribute to developing cardiomyocytes (Fig. 2**b**, Extended Data Fig. 3**b**). Consistent with these findings, we further discovered from a URD lineage inference analysis of previously published mouse embryonic scRNA-seq data^27^ that analogous LPM and LEM cells could be identified forming similar developmental trajectories contributing to developing cardiomyocytes (Extended Data Fig. 3**c-e**). Thus, our bioinformatic analyses support that cells with an extraembryonic signature (LEM) may contribute to the heart in a trajectory that is separate from that of the embryonic (LPM) lineage.

**Figure 2.**
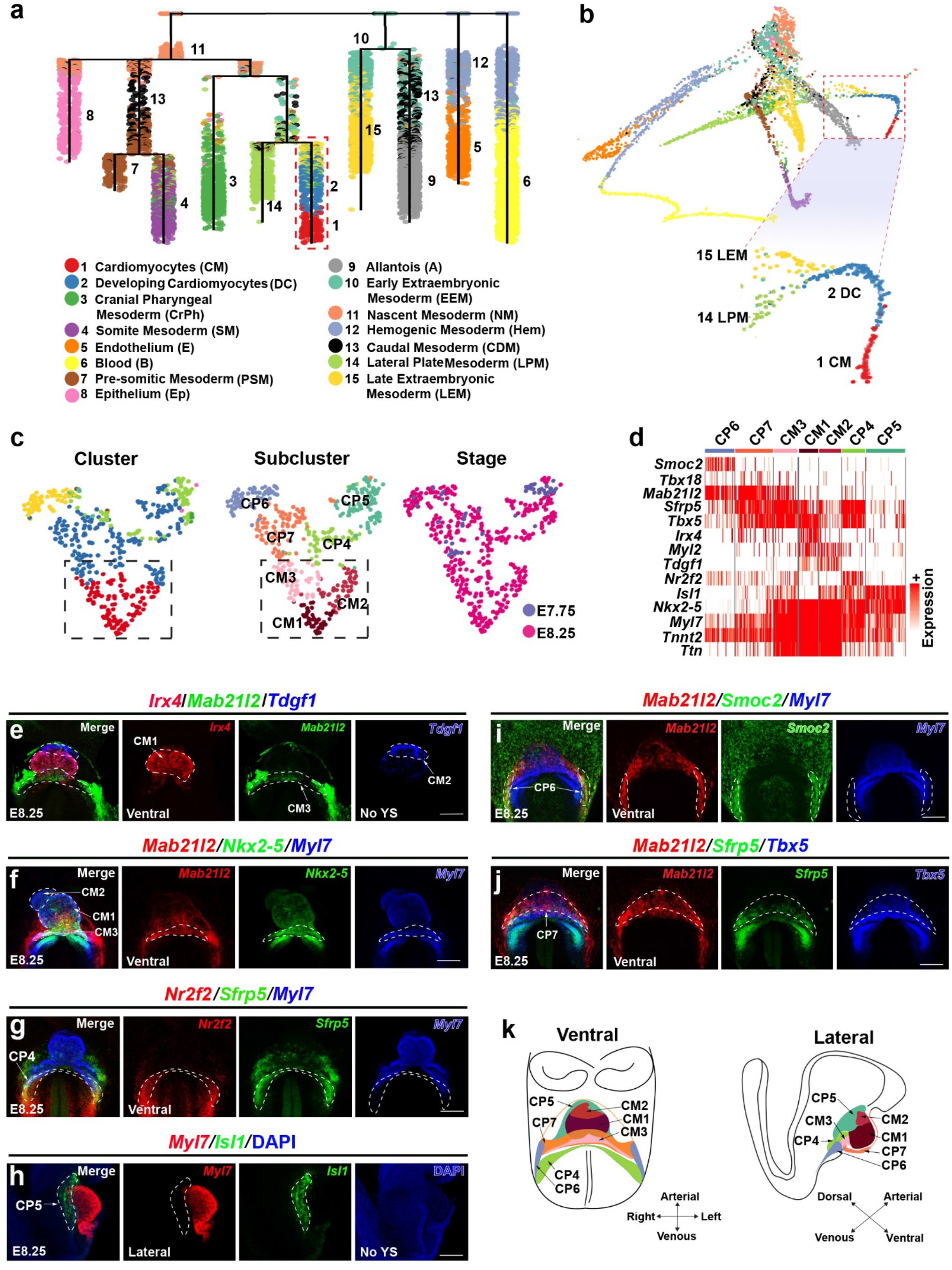
*Mesp1*-Cre scRNA-seq trajectory analysis reconstructs developmental cell lineage trees during mesoderm/heart organogenesis. **a, b**, URD inferred lineage tree, as displayed by (**a**) dendogram or (**b**) force-directed layout, reveals the developmental history of *Mesp1* mesoderm-derived organs. Red dashed box in **a**, **b** outlines cardiomyocyte branch, which is further magnified in **b.** The magnified cardiomyocyte branch shows that cardiomyocytes may derive from both late extraembryonic mesoderm (LEM) and lateral plate mesoderm (LPM) progenitor cells. **c**, tSNE layout of cells from only the cardiomyocyte branch (boxed area in **a**, **b**) reveals seven cardiac subclusters composing the cardiomyocyte branch including three distinct cardiomyocyte populations (CM1-3) and four specific cardiac progenitor cell-types (CP4-7). **d**, Heatmap of differentially expressed marker genes identifies each cardiac subcluster. **e-j**, RNAscope *in situ* hybridization (ISH) of representative marker genes for each cardiac subcluster cell population shows their location in E8.25 embryos. n = 3 per panel. Scale bars,100 μm. The extraembryonic tissue and part of the pericardium tissue were removed in **e**, **h** to show the underlying heart tube. **k**, Diagram illustrates the seven different cardiac subclusters in an E8.25 embryo.

### Multiple developmental pathways create distinct cardiomyocyte populations

Previous studies have reported the existence of distinct populations of cardiomyocytes during heart development which arise from distinct heart fields^2–4^. Thus, we investigated whether these specific cardiomyocyte populations could be detected as subclusters within our initially identified developing cardiomyocyte (DC)/cardiomyocyte (CM) clusters (Fig. 1**d**), and furthermore how LPM and LEM cells in our cardiomyocyte trajectories may specifically contribute to these subclusters (Fig. 2**b** - magnification). As a result, subclustering analysis of cells specifically comprising the initial developing cardiomyocyte/cardiomyocyte branches (Fig. 2**a, b** –boxed area, magnification: LEM, LPM, DC and CM) uncovered seven distinct sub-populations (Fig. 2**c, d**, Extended Data Fig. 4**a**, Supplementary Table 2). Three of these sub-clusters exhibited increased expression of cardiomyocyte sarcomeric genes such as *Ttn, Tnnt2, and Myl7* (Fig. 2**c, d**, Cardiomyocyte/CM1-3, Extended Data Fig. 4**a, b**), and correlated with the CM cluster (Fig. 2**c** - boxed area), whereas the other four sub-clusters displayed relatively low expression of these sarcomeric genes but high expression of cardiac progenitor (CP) markers such as *Isl1, Sfrp5, Tbx5* (Fig. 2**c, d**, Cardiac Progenitor/CP4-7, Extended Data Fig. 4**a, c**), and associated closely with the DC cluster and specific portions of LPM and LEM clusters (Fig. 2**c** - unboxed area). Differential gene marker analyses of the CM subclusters revealed that CM1, CM2 and CM3 cells displayed a combinatorial enrichment of *Irx4/Tbx5*, *Tdgf1*/*Isl1* and *Mab21l2/Tbx5*, respectively, and that CM1 cells exhibited increased expression of mature cardiomyocyte gene markers including *Actc1*, *Actn2, Myh6, Myh7* and *Myl1* (Extended Data Fig. 4**a, d**). Thus, these findings indicate that CM1 and CM2 subclusters may represent cardiomyocytes arising from the FHF and SHF^3–5,7,12,13,33–36^, whereas the developmental source of the CM3 subcluster remains to be identified. Additional gene marker analyses of CP subclusters support that cell types from some of these subclusters may represent cardiac progenitors for not only cardiomyocytes but also potentially other differentiated cardiac cell types (Fig. 2**d**, Extended Data Fig. 4**a, c**). For instance, CP6 and CP7 expressed genes that overlapped with those in not only CM3 cells but also proepicardial cells (*Upk3b*, *Ccbe1, Sfrp5, Mab21l2, Tbx18*^37–43^) (Extended Data Fig. 4**a**), suggesting that CP6/CP7 subclusters may contain progenitors for both CM3 and proepicardial cells.

To confirm the identity of potentially known subcluster cell types, annotate those that remain to be elucidated and further investigate their relationship during embryogenesis, we spatially mapped these cell types in E8.25 embryos when these cell types are present using RNAscope *in situ* hybridization (ISH) analysis of markers that are individually or combinatorially specific to these subclusters (Fig. 2**e-k**). Results from these studies revealed that *Irx4*, *Tdgf1* and *Mab21l2*, markers of CM1, CM2 and CM3 subclusters, respectively, were expressed in three distinct regions of the heart tube as labeled by *Myl7* and *Nkx2-5*: the middle segment (Primitive left ventricle/LV), arterial pole (Primitive outflow tract/OFT and right ventricle/RV), and venous pole of the heart tube, respectively (Fig. 2**e, f**), and thus indicate that CM1 and CM2 cells correspond to cardiomyocytes derived from the FHF and SHF, respectively^3–5,7,12,13,33–36^, whereas the source of progenitors giving rise to CM3 cardiomyocytes remains to be determined. Using a combination of genes that are differentially expressed in the CP subclusters, we further investigated the location of CP subcluster cell types during embryogenesis (Fig. 2**d, g-k**, Extended Data Fig. 4**a, c**). We discovered that the combined CP4 markers *Sfrp5* and *Nr2f2* were specifically expressed in regions posterior to the venous pole and contiguous with CM1 (Fig. 2**g**). The CP5 marker *Isl1* was enriched in regions anterior and dorsal to the arterial pole and contiguous with CM2 (Fig. 2**h**). The combined CP6 markers *Smoc2* and *Mab21l2* were expressed at the interface between the forming heart and extraembryonic tissues, near the ventral venous pole and contiguous with CM3 (Fig. 2**i**), and the combined CP7 markers *Sfrp5* and *Mab21l2* were located in a region connected to the ventral side of the venous pole and contiguous with CP6 (Fig. 2**j**). The adjacent locations of CP6 and CP7 and a large number of overlapping genes between them (*Cpa2, Mab21l2, Bmp4, Hand1*) (Fig. 2**d, i, j**, Extended Data Fig. 4**a, c**), suggest that CP6 and CP7 may be developmentally related.

Based on these subcluster analysis findings, we further investigated the developmental relationship of the CM1-3 subpopulations and specifically how LPM and LEM progenitors may contribute to them. To this end, we reconstructed our developmental trajectories (Fig. 3**a, b**, Extended Data Fig. 5) using the three CM subcluster populations CM1-3 (Fig. 2**c** - instead of the CM cluster from the initial tSNE cluster, Fig. 1**d**) as end points for our URD trajectory analysis^28^. As a result, the modified URD developmental trajectory tree created three new cardiomyocyte trajectory branches for each CM subcluster (Fig. 3**a, b**, box). The CM1 and CM2 trajectory branches, whose cells expressed genes associated with FHF (*Tbx5*) and SHF (*Isl1/Tdgf1*), respectively (Fig. 2**d**, Extended Data Fig. 4**a, b**), shared a common intraembryonic cellular origin associated with LPM and NM cells, whereas the CM3 lineage branch was distinct from the CM1 and CM2 branches and appeared to share an origin with early and late extraembryonic mesoderm (EEM and LEM) cells (Fig. 3**a, b**, Extended Data Fig. 5). Furthermore, the CM2 and cranial-pharyngeal (CrPh) branches expressed the SHF marker, *Isl1*, and appeared along a developmentally related trajectory consistent with previous studies of SHF development^5,6,8,9,11^ (Fig. **3a-c**).

**Figure 3.**
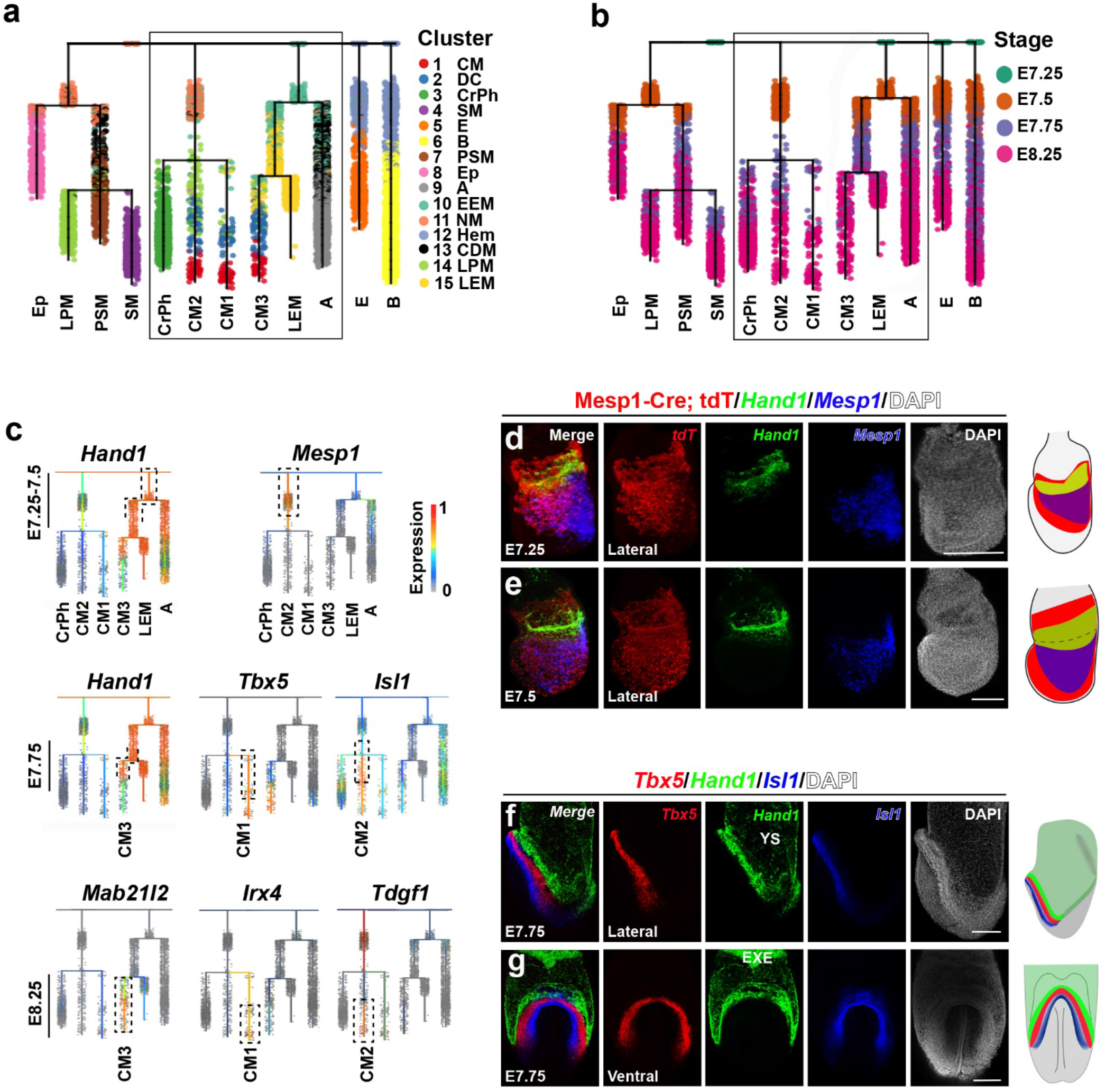
Distinct cardiomyocyte lineages derive from intra- and extra-embryonic related developmental origins. **a**, **b**, Reconstructed URD developmental cell lineage trees using the three distinct subclustered cardiomyocyte populations predict that CM1/CM2 and CM3 cardiomyocytes derive respectively from intra- and extra-embryonic related progenitor sources, as displayed by (**a**) cell type and (**b**) developmental stages. The cardiomyocyte-related branches of the URD developmental tree are outlined with box. **c**, Marker genes differentially expressed among the lineages for each cardiomyocyte subcluster are plotted on the URD cardiomyocyte-related branches. *Hand1* and *Mab21l2* mark early and late regions of the CM3 lineage, respectively. *Mesp1, Tbx5, Isl1, Irx4* and *Tdgf1* label different regions of the CM1 and CM2 lineage branches. **d**, **e**, RNAscope *in situ* hybridization (ISH) of *Mesp1* and *Hand1* was performed in (**d**) E7.25 and (**e**) E7.5 *Mesp1-Cre; Rosa26-tdT* embryos. The diagram illustrates both the gene expression pattern of *Hand1* and *Mesp1* and *Mesp1-Cre* lineage-traced cells in these embryos. **f**, **g**, RNAscope ISH of *Hand1, Tbx5*, and *Isl1* was performed in E7.75 embryos. The diagram illustrates the expression pattern of *Hand1*, *Tbx5*, and *Isl1* in these embryos. n = 3 per panel. Scale bars, 100 μm. EXE, Extraembryonic Ectoderm; YS, Yolk Sac.

### Interrogating transcriptional profiles of CM1-3 lineage branches uncovers distinct cell fate programs for each cardiomyocyte population

To identify gene programs that regulate the cell fate decisions creating these distinct cardiomyocyte lineages, we further interrogated the transcriptional profiles of cells along each of the cardiomyocyte developmental trajectories within the URD branching tree. To this end, we created a Random Forest model to classify and assign an importance score to transcription factors that may participate in directing cells to a specific daughter branch at each branch point examined in the URD tree^44^. These transcription factors were then ranked based on their importance score, and the top ten transcription factors that were predicted most likely to direct these branch point decisions were selected for each daughter branch (Extended data Fig. 6**a-c**). Expanding our analysis beyond transcription factors, we further identified the top twenty genes that were differentially expressed between daughter cells immediately after each branch point (Extended data Fig. 6**d-g**, Supplementary Table 3). These analyses revealed that *Hand1* appeared important for the initial branch point decision between embryonic NM and EEM (branch point 1) (Fig. 3**c**, Extended Data Fig. 6**a-d**), which coincides with the role of *Hand1* in extraembryonic mesoderm development^45,46^. Supporting previous cardiac developmental studies^3,5, 47–49^, we observed that the transcription factors *Tbx5, Isl1, Hand2 and Tbx1* exhibited differential expression and/or importance in FHF-related CM1, SHF-related CM2 and SHF-related CrPh cell-types at branch point 3 after their specification from intraembryonic NM/LPM cells (Figure 3**c**, Extended Data Fig. 6**a-c, f**). In the extraembryonic branch, *Cdx2/Cdx4* and *Tsc22d1* were reciprocally expressed at branch point 2 where allantois (A) and LEM cells arise from EEM cells (Extended Data Fig. 6**a, c, e**). Furthermore, *Mef2c*, *Id2* and *Cited2,* which were expressed in CM3 cells and have been implicated in cardiac development^50–54^, and *Hoxb6* and *Hand1,* which were expressed in the LEM cells, were predicted to play key roles in regulating cell fate decisions at branch point 4 (Extended Data Fig. 6**a-c, g**).

To further illuminate the dynamics of cell fate choices and corresponding differentiation states among these cardiomyocyte lineages, we examined genes differentially expressed in each cardiomyocyte lineage trajectory along a pseudotime from least to most differentiated conditions (Extended Data Fig. 7). These pseudotime analyses revealed at least three major differentiation states for each CM trajectory: an early, intermediate and late state (Extended Data Fig. 7**g-i**). Consistent with our branch point analyses (Extended Data Fig. 6), genes for the CM1 and CM2 early states were similar to each other but notably distinct from those for the CM3 early state; however, genes across these pseudotime analyses converged as each intermediate state cardiac progenitor differentiated into its corresponding late state cardiomyocyte population (Extended Data Fig. 7). In particular, *Mesp1* was expressed in the CM1 and CM2 early states but *Tbx5* and *Isl1* were reciprocally activated in these lineages at intermediate states (Fig. 3**c**, Extended Data Fig. 7**a, b, d, e, g, h**), suggesting that CM1 and CM2 may derive from a common developmental trajectory but *Tbx5* and *Isl1* may direct their specification in more distinct cardiomyocyte populations. On the other hand, *Hand1* and BMP signaling related genes *Bmp4* and *Msx2* were primarily expressed in CM3 early states, and *Mab21l2* and *Cpa2* were activated in CM3 intermediate states (Fig. 3**c**, Extended Data Fig. 7**c, f, i**). Finally, *Mef2c* and other sarcomeric genes (*Tnnt2*) were commonly expressed at the late state of CM1-3 lineages; however, some genes appeared to be specific for each CM population at this state including *Irx4* (CM1) and *Tdgf1* (CM2) (Extended Data Fig. 7**d-i**). Confirming these analyses, RNAscope ISH revealed that *Hand1* was expressed at the boundary of embryonic and extraembryonic tissues, whereas *Mesp1* was expressed in the proximal portion of the intraembryonic migrating mesoderm at E7.25 and E7.5 (Fig. 3**d, e**). Furthermore, *Hand1, Tbx5* and *Isl1* marked different locations in the crescent region at E7.75 where *Hand1* labeled a region anterior and lateral to the cardiac crescent, which was marked by *Tbx5*^3,34^, while *Isl1* labeled cells posterior and medial to the cardiac crescent as previously reported^5^ (Fig. 3**f, g**, Extended Data Fig. 8). Altogether, these bioinformatic and spatial gene expression analyses reveal a potentially unexplored developmental source of cardiomyocytes along the proximal extraembryonic-embryonic boundary that may be distinct from the previously described FHF and SHF progenitors.

### *Hand1* lineage tracing reveals an unexpected heart field that contributes to specific subsets of the first heart lineage and serosal mesothelial lineages

To examine and developmentally define this predicted extraembryonic-related developmental heart field, we employed an inducible Cre-recombinase genetic fate mapping strategy to lineage trace cells from this potential heart field during embryogenesis. Based on our interrogation of transcriptional profiles of cells along the CM URD trajectory branches, we discovered that *Hand1* was expressed in early extraembryonic-related CM3 progenitors but not CM1 and CM2 progenitors, thus identifying *Hand1* as a potential candidate gene to genetically label progenitors from the CM3 heart field (Fig. 3**c**). To further explore this possibility, we performed additional RNAscope ISH analyses to examine the expression of *Hand1* in the developing embryo and more specifically in these distinct cardiomyocyte progenitors. These studies revealed that *Hand1*+ mesoderm cells co-expressed *Mesp1* at the extraembryonic/embryonic boundary between E6.25 and E6.75 (Fig. 4**a**, Extended Data Fig. 9**a**) but downregulated *Mesp1* after E6.75 (Fig. 3**d, e**). Consistent with previous reports^55–57^, *Hand1* was expressed in the extraembryonic mesoderm at E7.75, E8.25 and pericardium at E8.25 but not in *Hcn4*+ or *Myl7*+ cardiomyocytes at any of these stages (Fig. 4**b, c**, Extended Data Fig. 9**b-e**). However, at E8.5 and E9.0, *Hand1* was expressed in a portion of cardiomyocytes in the LV and AVC as well as the pericardium and septum transversum (ST) (Extended Data Fig. 9**f, g**). Thus, the expression pattern of *Hand1* reveals a developmental time window (E6.25 - E8.25) in which the contributions of early gastrulating *Hand1+* CM3 progenitors to the heart can be investigated by *Hand1-CreERT2* lineage tracing.

**Figure 4.**
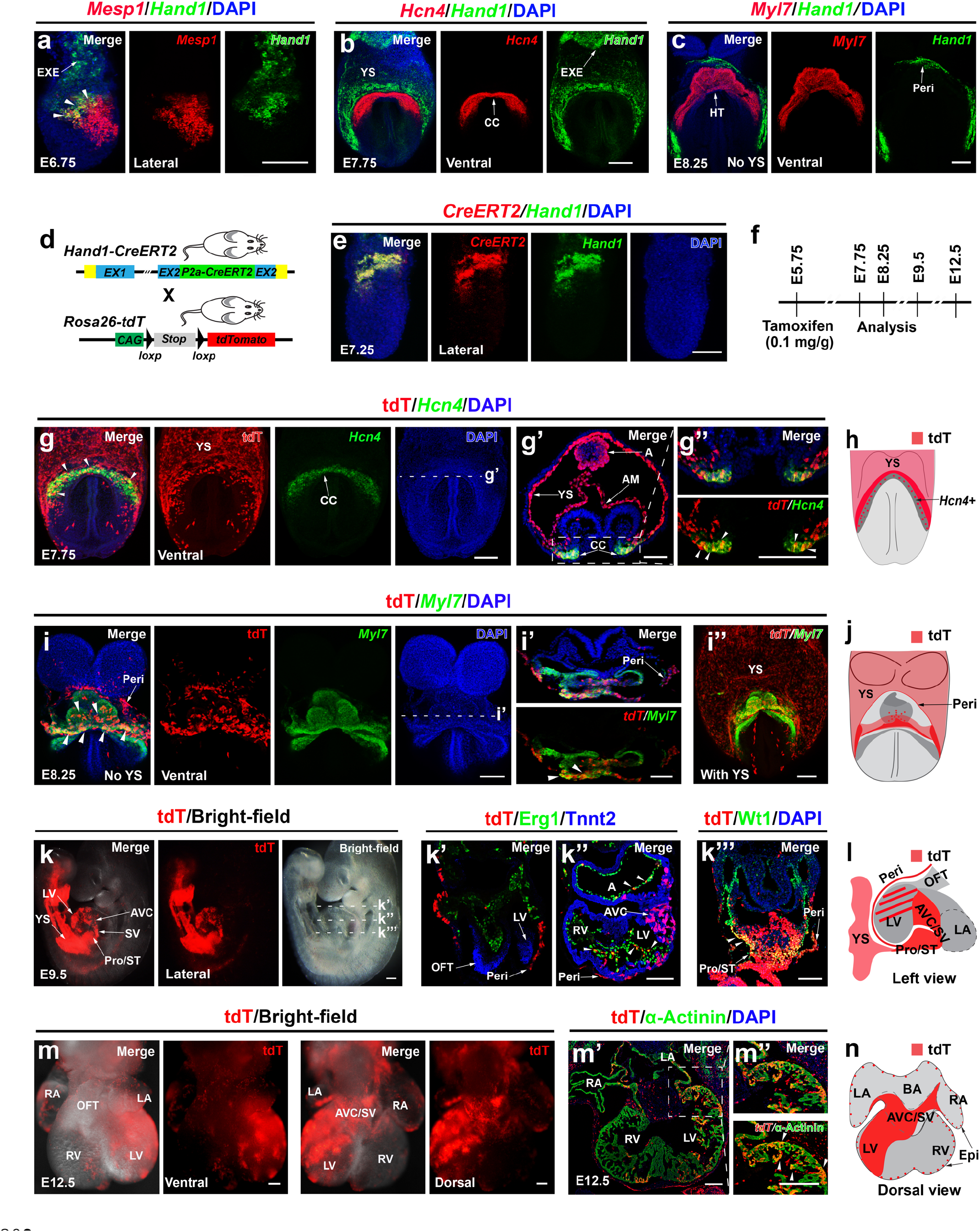
Lineage tracing studies reveal that early gastrulating *Hand1+* cells contribute to not only a distinct subpopulation of first heart lineage cardiomyocytes but also serosal mesothelial lineages (pericardial, epicardial cells) in the heart. **a-c**, RNAscope *in situ* hybridization (ISH) reveals that *Hand1* is expressed with (**a**) *Mesp1* at the embryonic and extraembryonic boundary in E6.75 embryos (arrowheads), (**b**) dorso-laterally around the cardiac crescent as detected by *Hcn4* at E7.75, and (**c**) in the pericardium which overlays the heart tube (HT) as detected by *Myl7* at E8.25. The yolk sac and part of the pericardium were removed in **c** to show the underlying heart tube. **d-n**, Lineage tracing studies using *Hand1-CreERT2* and *Rosa26-tdT* mice (shown in **d**) map the fate of early gastrulating *Hand1*+ cells. **e**, RNAscope ISH in *Hand1-CreERT2* embryos shows that expression of *CreERT2* precisely recapitulates the expression of *Hand1*. **f**, Schematic outlines the experimental strategy for *Hand1-CreERT2* genetic fate mapping studies shown in **g-n**. Tamoxifen was given at E5.75, and embryos were examined for *Hand1-CreERT2* genetically-labeled tdT+ cells at E7.75, E8.25, E9.5 and E12.5. **g-n**, RNAscope ISH and immunohistochemistry of whole mount and cross sections of these embryos reveal the contribution of *Hand1-CreERT2* genetically-labeled tdT+ cells at (**g**) E7.75, (**i**) E8.25, (**k**) E9.5 and (**m**) E12.5. **g’**, **i’**, **k’**, **k’’**, **k’’’**, Insets show transverse sections of **g**, **i**, **k** at indicated dashed lines, respectively. **m’**, Inset shows coronal section of **m. g’’**, **m’’**, Insets are magnification of **g’**, **m’** boxed area. Arrowheads point to tdT+ cells expressing (**g**, **g’’**) *Hcn4*, (**i**, **i’**) *Myl*7, (**k’’**) Erg1, (**k’’’**) Wt1 and (**m’’**) α-Actinin. **h**, **j**, **l**, **n**, Diagrams summarize the anatomical location of *Hand1-CreERT2* genetically-labeled tdT+ cells at the embryonic stages analyzed. n = 3 embryos for each stage. Scale bars, 100 μm. AM, Amnion; AVC, Atrioventricular Canal; BA, Base of the Atrium; CC, Cardiac Crescent; Epi, Epicardium; EXE, Extraembryonic Ectoderm; HT, Heart tube; LA, Left Atrium; LV, Left Ventricle; OFT, Outflow Tract; Peri, Pericardium; Pro, Proepicardium; RA, Right Atrium; RV, Right Ventricle; SV, Sinus Venosus; ST, Septum transversum; YS, Yolk Sac.

Accordingly, we generated a *Hand1-CreERT2* mouse by inserting a *P2a-CreERT2* cassette into the second exon of the *Hand1* gene (Fig. 4**d**, Extended Data Fig. 10**a, b**). RNAscope ISH studies confirmed that expression of *CreERT2* precisely recapitulated that of *Hand1* (Fig. 4**e**, Extended Data Fig. 10**c, d**). As a result, *Hand1-CreERT2* mice were bred with the *Rosa26-tdT* reporter mice to perform lineage tracing studies (Fig. 4**d**). Confirming the fidelity of the CreERT2 activity, Cre leakage was not observed in *Hand1-CreERT2; Rosa26-tdT* embryos without tamoxifen induction (Extended Data Fig. 10**e**). Because previous studies including our own (Extended Data Fig. 11) have shown that the half-life of tamoxifen in mice is ∼12 hours and persists over a ∼24–36 hour time period^58,59^, we studied *Hand1-CreERT2; Rosa26-tdT* embryos from pregnant mice given tamoxifen at E5.75 (Fig. 4**f**) to avoid the possibility that a small amount of residual tamoxifen would activate *CreERT2* in differentiated cardiomyocytes expressing *Hand1* at E8.5. Consistent with our CM trajectory branches (Fig. 3**c**), examination of these genetically-labeled embryos at E7.75, E8.25, E8.5, E9.5 and E12.5 revealed that *Hand1* lineage-traced cells contributed to not only extraembryonic tissue but also the heart (Fig. 4**g-n**, Extended Data Fig. 12). Within extraembryonic tissues, which were tdT-labeled throughout the yolk sac across all examined stages (Fig. 4**g-l**, Extended Data Fig. 12**b**), *Hand1* lineage-traced tdT+ cells produced Pecam+ endothelial cells, α-SMA+ smooth muscle cells and Pdgfrβ+ mesothelial cells (Extended Data Fig. 12**e, f**). On the other hand, *Hand1* lineage-traced tdT+ cells contributed to the developing embryo in a more spatial and temporal restricted manner (Fig. 4**g-n**). Specifically, *Hand1* lineage-traced tdT+ cells supplied *Hcn4+* cardiomyocytes in the cardiac crescent at E7.75 and then cardiomyocytes (*Myl7*+) on the ventral side of the venous pole and medial regions of the heart tube at E8.25 (Fig. 4**g-j**). At later stages (E8.5 - E12.5), tdT+ cardiomyocytes were increasingly restricted spatially to the primitive AVC region and LV at E8.5 and then further to the AVC/sinus venosus (SV), dorsolateral LV and atrial regions of the heart at E12.5 (Fig. 4**k-n**, Extended Data Fig. 12**b, d**). Furthermore, tdT+ cells appeared in non-myocardial heart tissue including the pericardium, proepicardium/ST, epicardium, and occasionally endocardium from E8.25 - E12.5 (Fig. 4**i-n**, Extended Data Fig. 12**a-d**). Supporting these lineage studies, the CM3 URD tree branch, which included the CP6 and CP7 subclusters, comprised cells that express AVC markers (*Msx1/2, Twist1, Tbx2*^42,60–62^), as well as proepicardial/pericardial markers (*Upk3b, Ccbe1, Sfrp5, Mab21l2, Tbx18*^37–43^) (Extended Data Fig. 4**a**). Finally, to confirm these findings and investigate additional early gastrulating *Hand1+* CM3 progenitors that may not have been labeled at E5.75, we examined *Hand1-CreERT2; Rosa26-tdT* embryos from pregnant mice given tamoxifen at E6.25 (Extended Data Fig. 13). In addition to displaying similar results to those observed in E5.75 tamoxifen-induced *Hand1-CreERT2; Rosa26-tdT* embryos (Extended Data Fig. 13**a-f** compared to Fig. 4**g-n**), *Hand1* lineage-traced tdT+ cells genetically-labeled at E6.25 also contributed cardiomyocytes as well as all epicardial-derived cell-types including fibroblasts and vascular support cells to E17.5 embryos (Extended Data Fig. 13**g**). Altogether, these data suggest that at early gastrulation stages, *Hand1* marks a progenitor population that gives rise not only to cardiomyocytes within the AVC and LV prior to the time when *Hand1* is actively expressed in differentiated CMs, but also to pericardial, epicardial and extraembryonic derived mesoderm cell types. As these *Hand1+* cardiomyocyte progenitors specifically contribute to cardiomyocytes within the developing atrioventricular canal and dorsolateral regions of the LV, they likely represent a distinct subset of the reported first heart lineage cardiomyocytes^2^, suggesting that the FHF is not a single heart field, but is rather composed of at least two distinct heart fields, one of which, identified here, is marked by *Hand1*.

### Genetic clonal analysis reveals the multipotentiality of *Hand1+* cardiac precursor cells

Our lineage tracing results reveal that *Hand1+* progenitors in the early gastrulating embryo give rise to multiple distinct cell types in extraembryonic tissue, pericardium, endocardium, epicardium as well as the dorsal LV and AVC myocardium in the developing heart (Fig. 4, Extended Data Fig. 12, 13). These findings may reflect the presence of distinct *Hand1+* precursor cells that produce individual cell types, or multipotential *Hand1+* precursor cells which can differentiate into different combinations of cell types. To examine the lineage potential of single *Hand1*-expressing cells during early gastrulation, we crossed *Hand1-CreERT2* mice with the *Rosa26-Confetti* multicolor reporter mice^63^ to genetically fate map early *Hand1+* individual clones expressing a specific fluorescent protein following low dose tamoxifen treatment at E6.75 or E7.25 (Fig. 5**a**). To this end, we discovered that 0.005 mg/g of tamoxifen was the minimum effective dose at E6.75 or E7.25 that reliably leads to clonal events in examined embryos (Extended Data Fig. 14**a, b**). This dose resulted in recombination in only 27% of embryos (n = 175/640), which was less than the expected *Hand1-CreERT2*; *Rosa26-confetti* genotype positivity rate (50%) (Extended Data Fig. 14**b**). Among these embryos, bi-color embryos occurred in the highest proportion followed by uni-color and tri-color embryos (Fig. 5**b**), and the observed frequency of each Brainbow color (YFP, RFP, CFP and nGFP) was consistent with those from previous reports^4^ (Extended Data Fig. 14**c**). Further examining all labeled embryos at E9.5 revealed that ∼73% (242/330) of the clones were present in only the extraembryonic tissue; ∼26% (86/330) were located in both extraembryonic and cardiac tissue; and two clones (∼1%) contributed to only cardiac tissue (Extended Data Fig. 14**d**). The distribution of these clones combined together were consistent with the distribution of genetically-labeled *Hand1-CreERT2*; *Rosa26-tdT* cells at a similar stage (Fig. 4**k, l**, compared to Fig. 5**c-e)**. Supporting the multipotentiality of early *Hand1+* progenitors during early gastrulation, uni-color embryos, which were most likely due to a single recombination event^4^, exhibited clones that fluorescently-labeled both extraembryonic and cardiac tissues including the proepicardium/ST, pericardium, LV and AVC at E9.5 (Fig. 5**c-e**). Additional immunofluorescence studies revealed that these *Hand1+* clones specifically contributed to not only α-Actinin+ cardiomyocytes in the AVC and LV but also Wt1+ proepicardial and pericardial cells in the developing embryo (Fig. 5**c**, Extended Data Fig. 14**e-h**). However, consistent with the relatively few endocardial cells genetically labeled in *Hand1-CreERT2; Rosa26-tdT* embryos (Fig. 4**k’’**, Extended Data Fig. 12**d’**), *Hand1+* clones were not observed in the endocardium. Finally, *Hand1+* clones supplied α-SMA+ smooth muscle cells and Pdgfrβ+ mesothelial cells to the extraembryonic mesoderm (Extended Data Fig. 14**i, j**).

**Figure 5.**
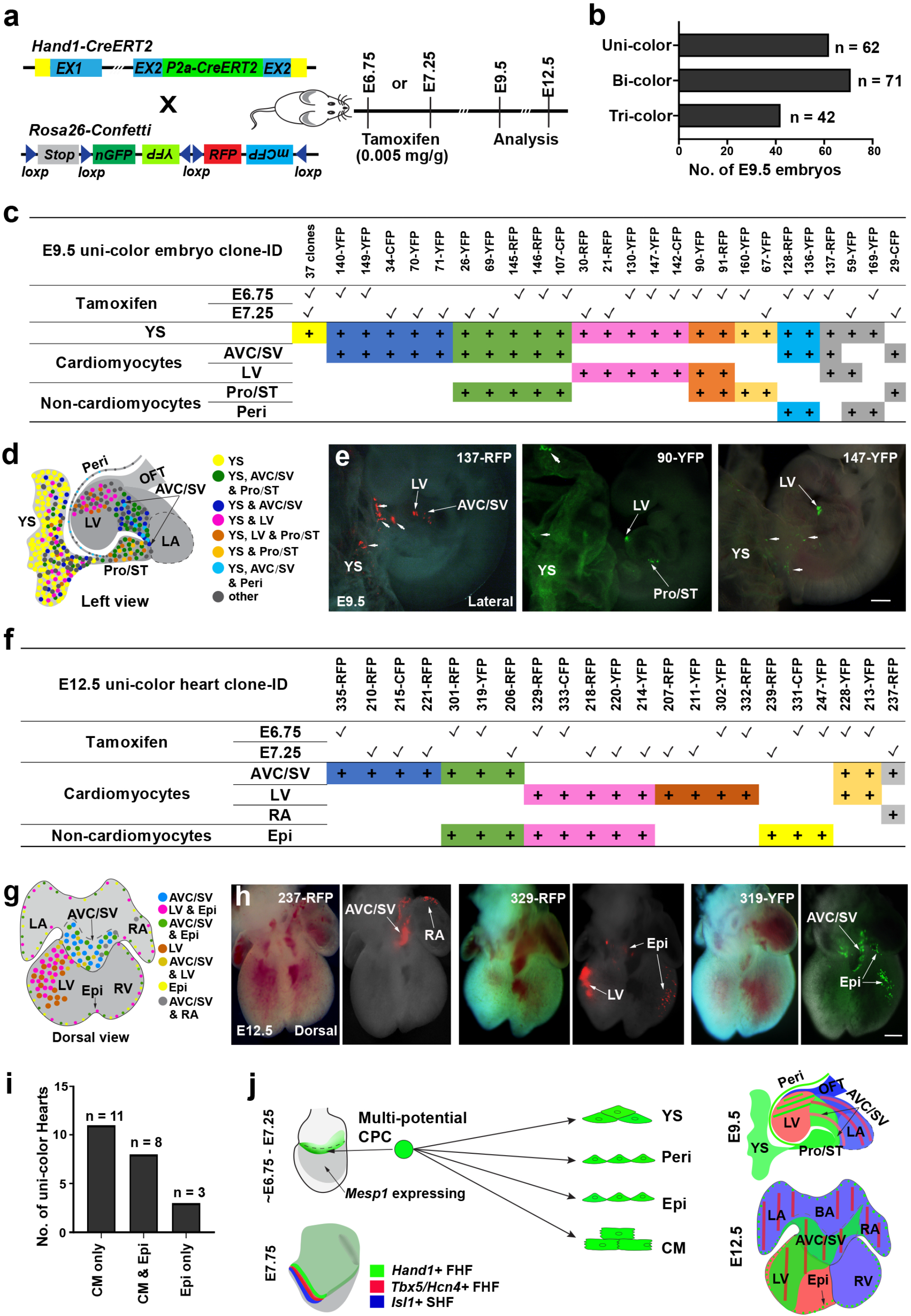
Clonal analysis reveals multipotentiality in early *Hand1+* progenitors. **a**, Schematic outlines experimental strategy for *Hand1-CreERT2; Rosa26-Confetti* clonal analyses. **b**, Bar graph displays the number of uni-color, bi-color or tri-color *Hand1-CreERT2; Rosa26-Confetti* embryos at E9.5. **c**, **f**, Clonal analyses of uni-color (**c**) E9.5 and (**f**) E12.5 embryos reveal that individual *Hand1-CreERT2; Rosa26-Confetti* clones labeled at E6.75 or E7.25 have the capacity to generate multiple cell types that can contribute to the yolk sac and/or heart. **d**, **g**, Diagram summarizes the contribution of *Hand1-CreERT2; Rosa26-Confetti* genetically-labeled clones in the heart and yolk sac at (**d**) E9.5 and in the heart at (**g**) E12.5. **e**, **h**, Representative (**e**) E9.5 and (**h**) E12.5 uni-color embryos show individual *Hand1-CreERT2; Rosa26-Confetti* genetically-labeled clones contributing to different combinations of tissues and cell types: (**e**) AVC/SV, LV and YS (clone #137-RFP); LV, Pro/ST and YS (clone #90-YFP); LV and YS (clone #147-YFP); (**h**) AVC/SV and RA (clone # 237-RFP); LV and Epi (clone # 329-RFP); the AVC/SV and Epi (clone # 319-YFP). Arrowheads point to yolk sac cells in **e**. Scale bars, 200 µm. **i**, Bar graph displays the number of uni-color E12.5 hearts with clones contributing to cardiomyocytes only, epicardial cells and cardiomyocytes, or only epicardial cells. **j**, Model summarizes the multipotentiality of *Hand1+* cardiac progenitor cells (CPC) between E6.75 - E7.25 in relation to the contribution of reported FHF/SHF progenitors. AVC, Atrioventricular Canal; BA, Base of the Atrium; CM, cardiomyocytes; Epi, Epicardium; LA, Left Atrium; LV, Left Ventricle; OFT, Outflow Tract; Peri, Pericardium; Pro, Proepicardium; RA, Right Atrium; RV, Right Ventricle; SV, Sinus Venosus; ST, Septum Transversum; YS, Yolk Sac.

To further substantiate that clones in both the yolk sac and heart derive from a single recombination event, we examined additional embryos containing bi- and tri-color clones. To ensure that these clones resulted from single recombination events, we employed a rigorous statistical analysis of the number cells in each clone. To this end, we counted the cells in each clone (an individual color) in either cardiac tissues or both extraembryonic and cardiac tissues, and modeled these cell counts with a mixture of two Gaussian distributions^64^: one for the cell count that would be expected for a single recombination event, and the other for the cell count that would be expected for two or more recombination events (Extended Data Fig.14**m-p**). Based on this model, we found that 53 out of 88 observed clones labeled at E6.75 or E7.25 corresponded to single clonal events in both cell count analyses (Extended Data Fig. 14**m-p**). Additional analyses of these consensus single clonal events revealed that the majority of single clones in the heart contributed to two or three distinct lineages, including various combinations of extraembryonic mesoderm, pericardium, proepicardium/ST, and AVC or LV myocardium (Extended Data Fig. 14**n, p**), thus supporting the multipotentiality of *Hand1*+ progenitor cells.

To further investigate the clonal relationships among specific *Hand1*+ progenitor-derived cardiac cell types and their location in later stage hearts, we examined E6.75 or E7.25 tamoxifen-induced *Hand1-CreERT2*; *Rosa26-Confetti* clones at E12.5 when most cardiac structures and cell types have been determined. Only uni-color hearts were analyzed as these hearts were most likely to be derived from a single recombination event^4^. Consistent with the E9.5 clonal analysis (Fig. 5**c**), clones marking the epicardium also labeled cardiomyocytes in the AVC or LV at E12.5 (Fig. 5**f-i**, Extended Data Fig. 14**k**), thus supporting that multipotential *Hand1+* cardiac progenitor cells can give rise to both cardiomyocytes and non-cardiomyocytes. Altogether, these results reveal the existence of multipotential *Hand1* cardiac progenitors in the early ingressing mesoderm that can give rise to extraembryonic mesoderm, mesothelial lineages (epicardium and pericardium) and LV and AVC myocardium (Fig. 5**j**).

## Discussion

Overall, our transcriptional and developmental interrogation of *Mesp1*-lineages at single-cell resolution has illuminated the intricacies of building complex organs/tissues derived from the mesoderm. Our single-cell transcriptomic studies reveal not only well-established but also previously unappreciated developmental sources for key cell lineages creating both intra- and extra-embryonic organs/tissues. Similar to previous studies for gut endoderm^65^, our trajectory analysis of our developing mesoderm single cell data has uncovered a close developmental relationship between intra- and extra-embryonic derived organs/tissues including unexpectedly a distinct developmental lineage of the heart that is related to those contributing to specific extraembryonic structures. Utilizing a combination of genetic fate-mapping and clonal analyses, we not only confirm this developmental cardiac-extraembryonic tissue connection but also delineate the progenitors creating these lineages and their specific contributions to the developing heart and extraembryonic structures (Fig. 5**j**).

Highlighting the complexity of organogenesis, we show how similar cell types, such as cardiomyocytes, can derive from multiple developmental origins/progenitors that have potential to contribute not only to other cell types but also to multiple organs/tissue structures. In particular, further single-cell subcluster analyses of isolated cardiomyocyte transcriptomic profiles identified at least three distinct myocardial heart lineages including a heart lineage whose progenitor shares a gene signature with extraembryonic mesodermal progenitors including *Hand1*. Trajectory analyses predicted that two of these heart lineages derive from a common embryonic source prior to E7.25, with marker expression at E8.25 suggesting their correspondence to first and second heart lineages^1^, whereas the *Hand1*+ extraembryonic-related heart lineage originates from a distinct developmental source that downregulates *Mesp1* prior to E7.25 and gives rise to myocardial lineages. Further expression analyses revealed that at early gastrula stages, *Hand1*+ progenitors reside at the intra-/extra-embryonic boundary, with genetic fate mapping demonstrating that *Hand1*+ progenitors specifically contribute to myocardial cells localized to the dorsal regions of the LV and AVC at E12.5. As myocardial lineages contributing to the LV have previously been defined as first heart lineages deriving from the FHF^2^, our results support that this *Hand1+* cardiac progenitor field represents a distinct subset of the FHF, thus revealing that the FHF consists of at least two distinct progenitor fields. Notably, these *Hand1*+ FHF subpopulation findings are consistent with a mathematically-inferred myocardial lineage model from previous retrospective clonal analyses^2–4^, thus further supporting our findings.

One limitation for understanding the full lineage potential of the FHF or SHF from retrospective clonal studies^2^, is that, because of experimental design, only myocardial clones can be studied. However, when *Isl1* was identified as a marker of the SHF, studies with *Isl1-Cre* or inducible *Isl1-CreERT2* revealed that the SHF produces both myocardial lineages as well as multiple other cardiac lineages^5,8,66,67^. Here, utilizing *Hand1-CreERT2* in concert with a confetti clonal indicator^63^, we uncovered an unsuspected multipotentiality of the *Hand1* FHF in which cells within the *Hand1* FHF can give rise not only to a specific subset of myocardial lineages within the first heart lineage, but also to extraembryonic mesoderm, septum transversum/epicardial, and pericardial cells. Thus, our results reveal that myocardial cells of the AVC and LV (particularly dorsal regions) and extraembryonic mesodermal and serosal mesothelial cells have a closer lineage relationship than previously expected, while also addressing the elusive embryonic origins of the proepicardium/epicardium, which contributes essential vascular support cells and cardiac fibroblasts to the heart.

The existence of a progenitor population that gives rise to cells both within extraembryonic and intraembryonic tissues provides a further example of blurred boundaries between extraembryonic and intraembryonic tissues, as seen by migration of extraembryonic hematopoietic progenitors to intraembryonic sites^68^, and intercalation of extraembryonic and intraembryonic endoderm during gut formation^69^. Additionally, these findings may also account for previous observations that, under certain *in vitro* conditions, epicardial progenitors can adopt cardiomyocyte cell fates^70^, and that loss of an endothelial/hematopoietic transcription factor, Scl, can result in transdifferentiation of yolk sac hematopoietic cells to beating cardiomyocytes^71^. The close developmental relationship between mesothelial lineages of both extraembryonic and intraembryonic tissues, and *Hand1+* FHF-derived cardiac lineages, coupled to the high plasticity of mesothelial cells^72–74^, suggests the possibility of transforming extraembryonic and serosal mesothelial tissues into cardiomyocytes to treat heart failure in the future.

As *Hand1* marks a subset of the FHF in the early gastrula embryo, and these progenitors have multipotentiality, the specific role of *Hand1* in early specification of FHF progenitors will be of great interest to examine in future studies. Global knockout of *Hand1* results in embryonic lethality at approximately E8.5, and mutant embryos exhibit placental, yolk sac and heart defects^45,46^. As placenta and yolk sac defects can secondarily impact the heart, direct requirements for *Hand1* in early heart progenitors remains unclear. Although some experiments, including cardiac-specific conditional knockout and tetraploid rescue studies^45,46^, confirmed heart defects in *Hand1* mutant embryos, these studies could not rule out requirements for *Hand1* in differentiated cardiomyocytes, rather than undifferentiated progenitors. However, our findings suggest that these heart defects may be due to abnormalities in undifferentiated progenitors which can give rise to cardiomyocytes. Because these progenitors can also contribute to extraembryonic tissue, they also raise the possibility that congenital heart diseases thought to be caused by placental anomalies^75^ may be due to perturbations of complex interplays between genetic pathways shared by extraembryonic and cardiac lineages.

Overall, our studies reveal that there are distinct subsets of the FHF that contribute to specific corresponding subpopulations of first myocardial lineages^2^, and that *Hand1*-FHF progenitors are multipotential, giving rise to multiple cell lineages, including cardiovascular lineages within the heart and extraembryonic cell types within the yolk sac. The cardiovascular multipotentiality of FHF progenitors, as previously seen for the SHF, may further reflect the evolution of the cardiovascular system, thus highlighting the overall complexity of how diverse cell types are created to build and organize functional organs/tissues.

## Acknowledgements

We thank Jianlin Zhang and Mi Tran for mouse care, and Evans and Chi lab members for comments on the manuscript. Various experiments were conducted with the assistance, expertise, and support of the following University of California San Diego (UCSD) core facilities including the: Institute for Genomic Medicine Core, Mouse Transgenic Core and Histology and Immunohistochemistry Core. This work was supported in part by grants from the NIH to S.M.E., N.C.C.; CIRM to T. I., N.C.C.; AHA to J.B.

## Author Contributions

Q.Z., D.C., P.C., S.M.E., J.B. and N.C.C. conceived the project and the design of the experimental strategy. Q.Z., J.B. carried out experimental studies including *in situ* hybridization study, lineage tracing and clonal analysis. D.C. performed bioinformatic analysis. J.B. carried out scRNA-seq experimental studies. F.Z. generated *Hand1-CreERT2* knock in mice line. T. I provided bioinformatics analysis guidance. Q.Z., D.C., S.M.E., J.B. and N.C.C. prepared the manuscript.

## Competing interests

The authors declare no competing interests.

Reprints and permission information is available at www.nature.com/reprints.

---

**Extended Data Figure 1.**
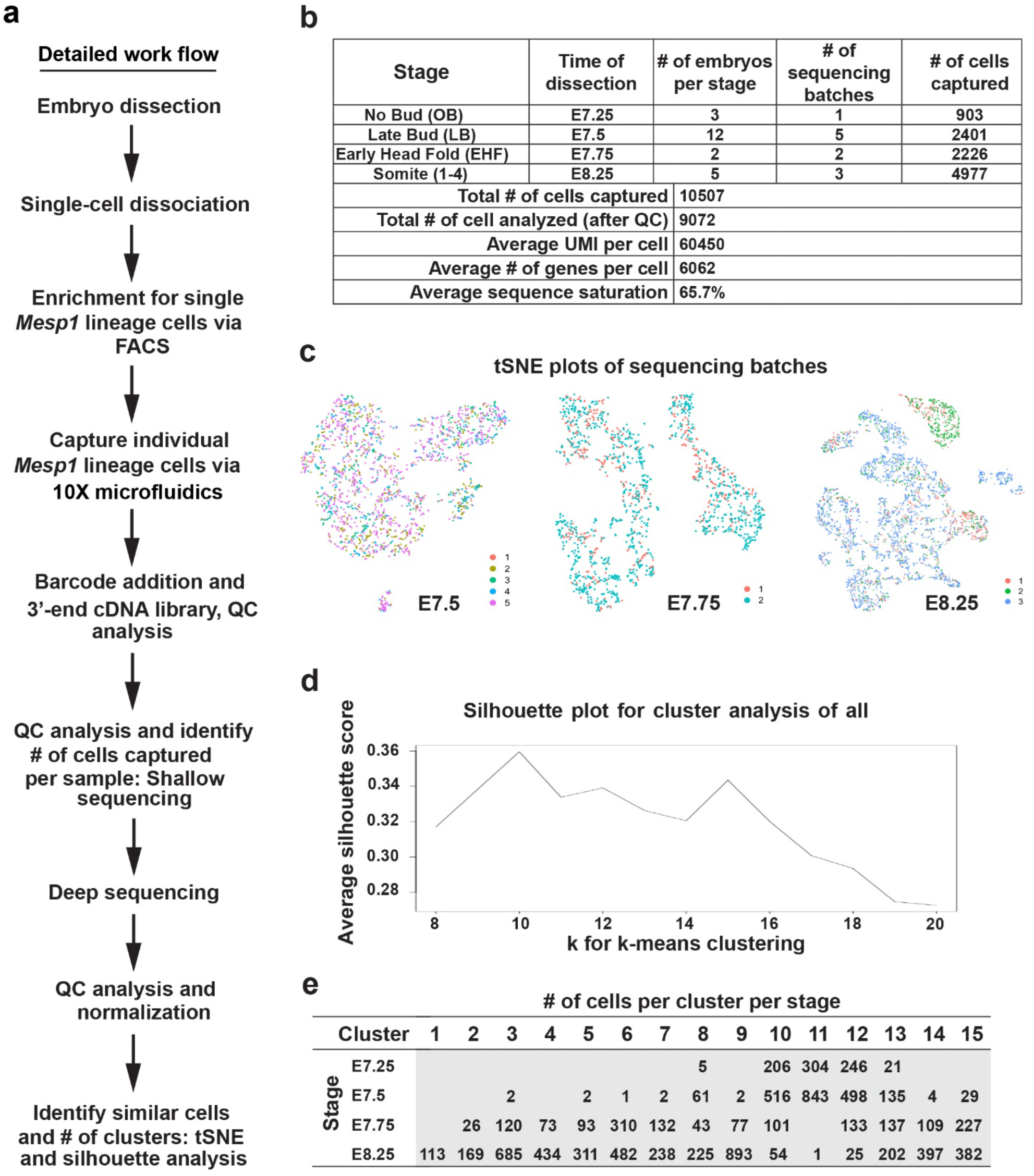
*Mesp1*-Cre scRNA-seq experimental details show the sequencing, quality control and analysis parameters used on single mesoderm cells. **a**, Diagram outlines details of the scRNA-seq workflow. **b**, Table shows the number of embryos and cells harvested and studied at each experimental condition. **c**, tSNE plots of sequencing experiments show that scRNA-seq datasets are well mixed at each developmental stage with no appreciable batch effects. **d**, Average silhouette score shows how the number of clusters for scRNA-seq data was determined. **e**, Table shows the number of cells sequenced and analyzed in each identified cluster at specific stages.

**Extended Data Figure 2.**
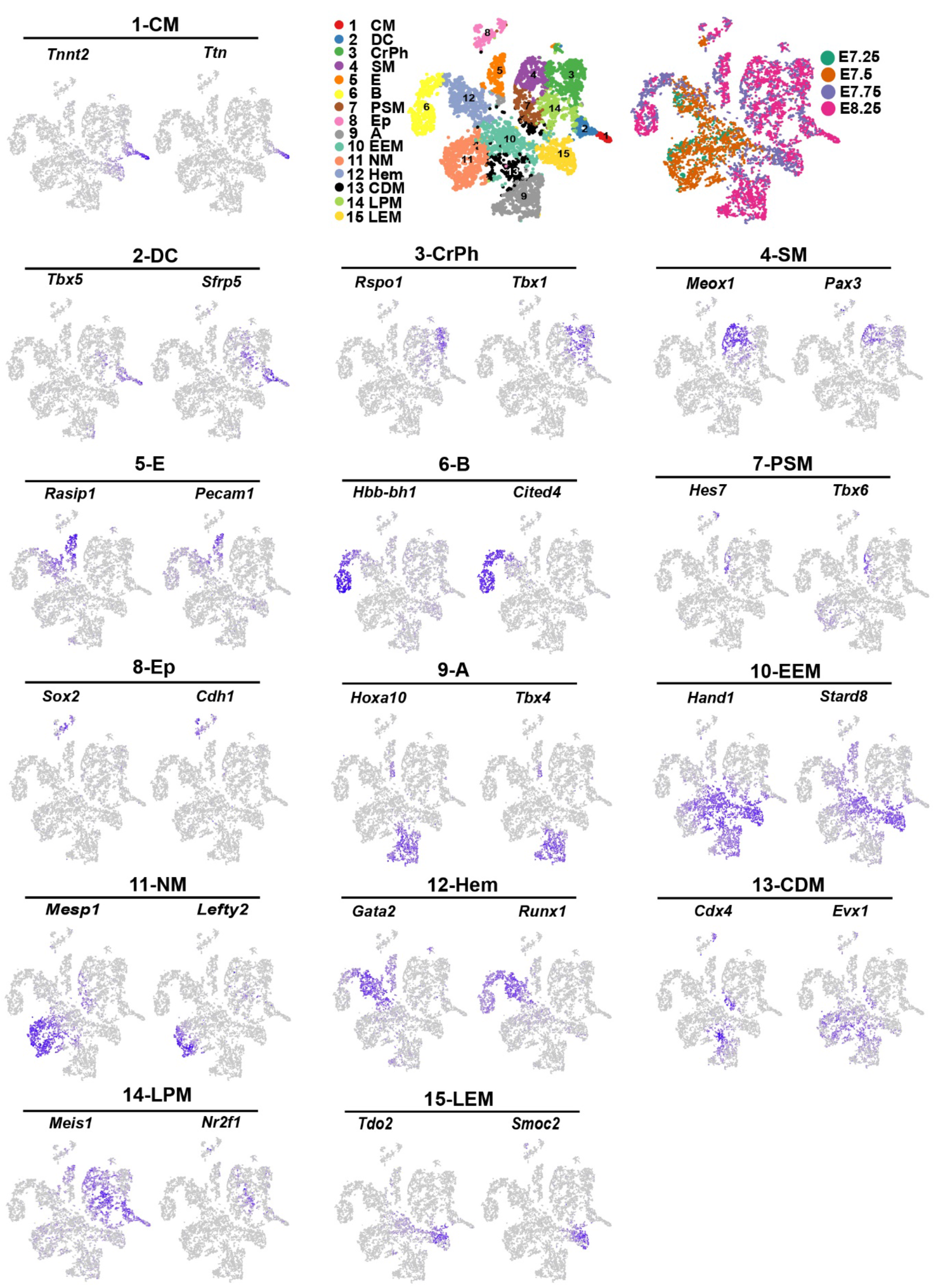
The expression of gene markers specific to identified clusters is visualized in tSNE plots of *Mesp1*-Cre scRNA-seq data. The expression of two representative markers for each cluster are displayed on the single-cell tSNE layout of all developmental stages.

**Extended Data Figure 3.**
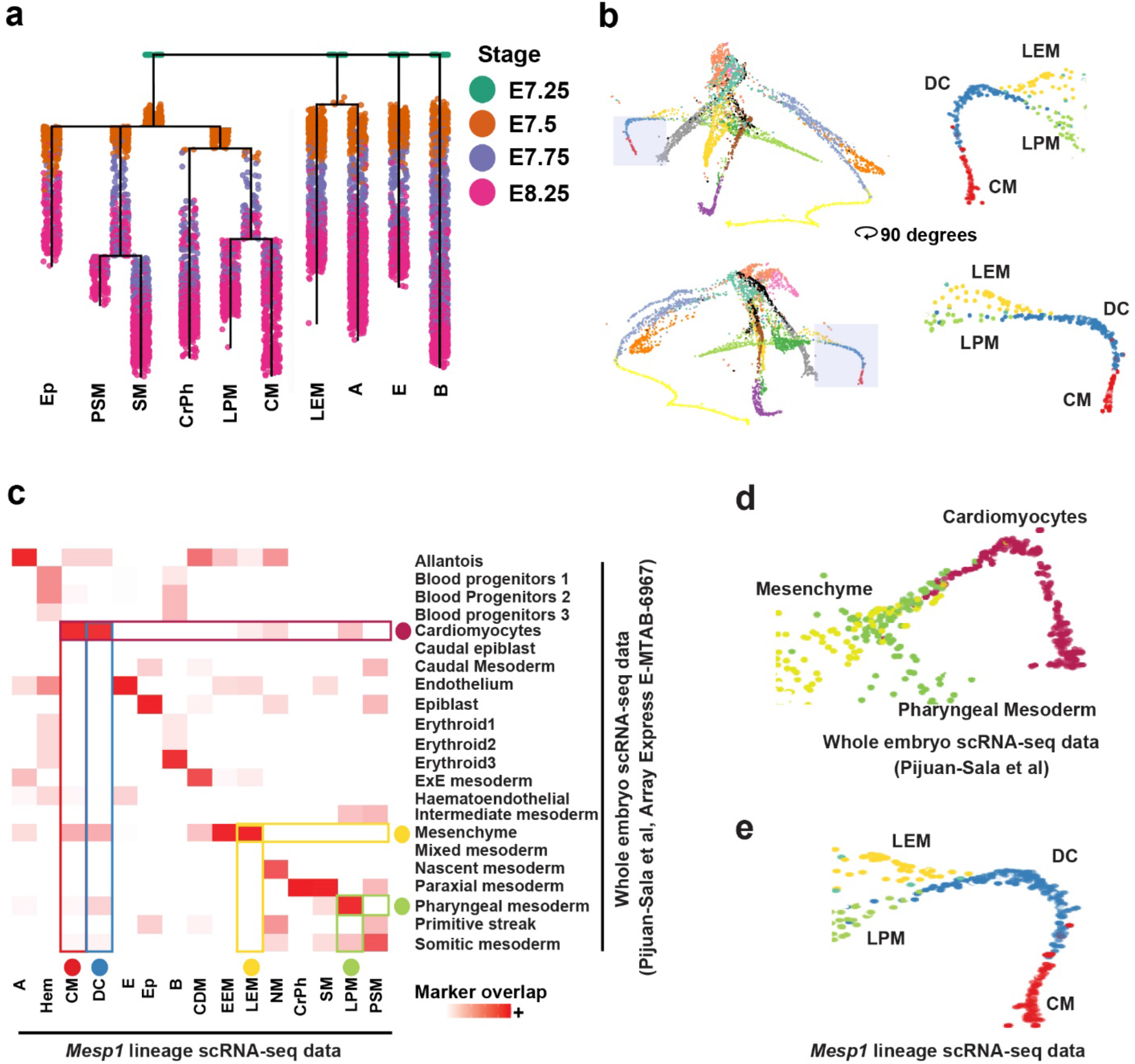
Trajectory analysis in an independent developing mouse embryo scRNA-seq dataset predicts similar intra- and extra-embryonic related developmental sources of cardiac progenitors. **a**, Developmental stage of each RNA-sequenced *Mesp1*-Cre labeled cell was projected onto *Mesp1*-Cre URD developmental lineage tree reconstructed in Figure. 2**a**. **b**, 3D rotations of the force-directed *Mesp1*-Cre URD layout provide different views of late extraembryonic mesoderm (LEM) and lateral plate mesoderm (LPM) trajectories converging to form the cardiomyocyte branch. Shaded branches are magnified and shown to the right. **c**, Heatmap of identified cell-type clusters from *Mesp1*-Cre and independent mouse embryonic scRNA-seq datasets^27^ reveals corresponding analogous cell-type populations between the datasets. **d**, **e**, An URD-derived force-directed layout of these independent scRNA-seq datasets shows how their analogous cell types exhibit similar converging trajectories that create cardiomyocytes.

**Extended Data Figure 4.**
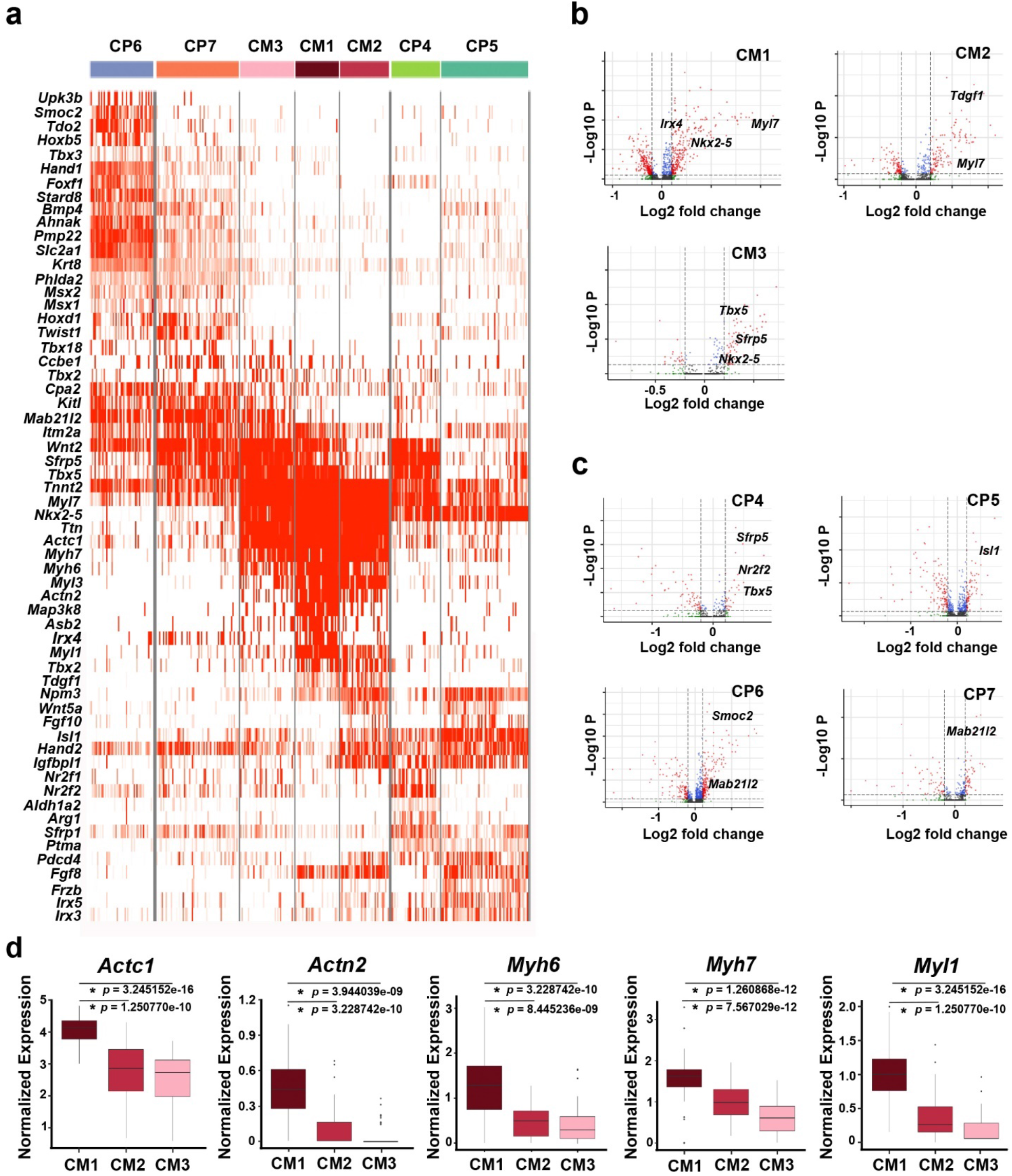
Differential gene expression analysis reveals cardiac subtype identities and transcriptional profiles. **a**, An extended heatmap of differentially expressed genes among cardiac subclusters reveals distinct transcriptional profiles for each developmental cardiac subcluster cell type derived in Figure 2**c. b, c**, Volcano plots summarize the differential expression of each cardiac subcluster (**b**, cardiomyocyte subclusters and **c**, cardiac progenitor subclusters) compared to all other cardiac subclusters. **d**, Box plots show that CM1 cells display higher expression of *Actc1, Actn2, Myh6, Myh7* and *Myl1* than that observed in CM2 and CM3. Median, 25^th^ and 75^th^ quartile, and extreme values within 1.5 times the interquartile range are indicated by the center line, bottom and top of the box, and ends of whiskers, respectively. Outliers outside 1.5 times the interquartile range appear as individual points. \**p* < 0.01 by Wilcoxon rank sum test.

**Extended Data Figure 5.**
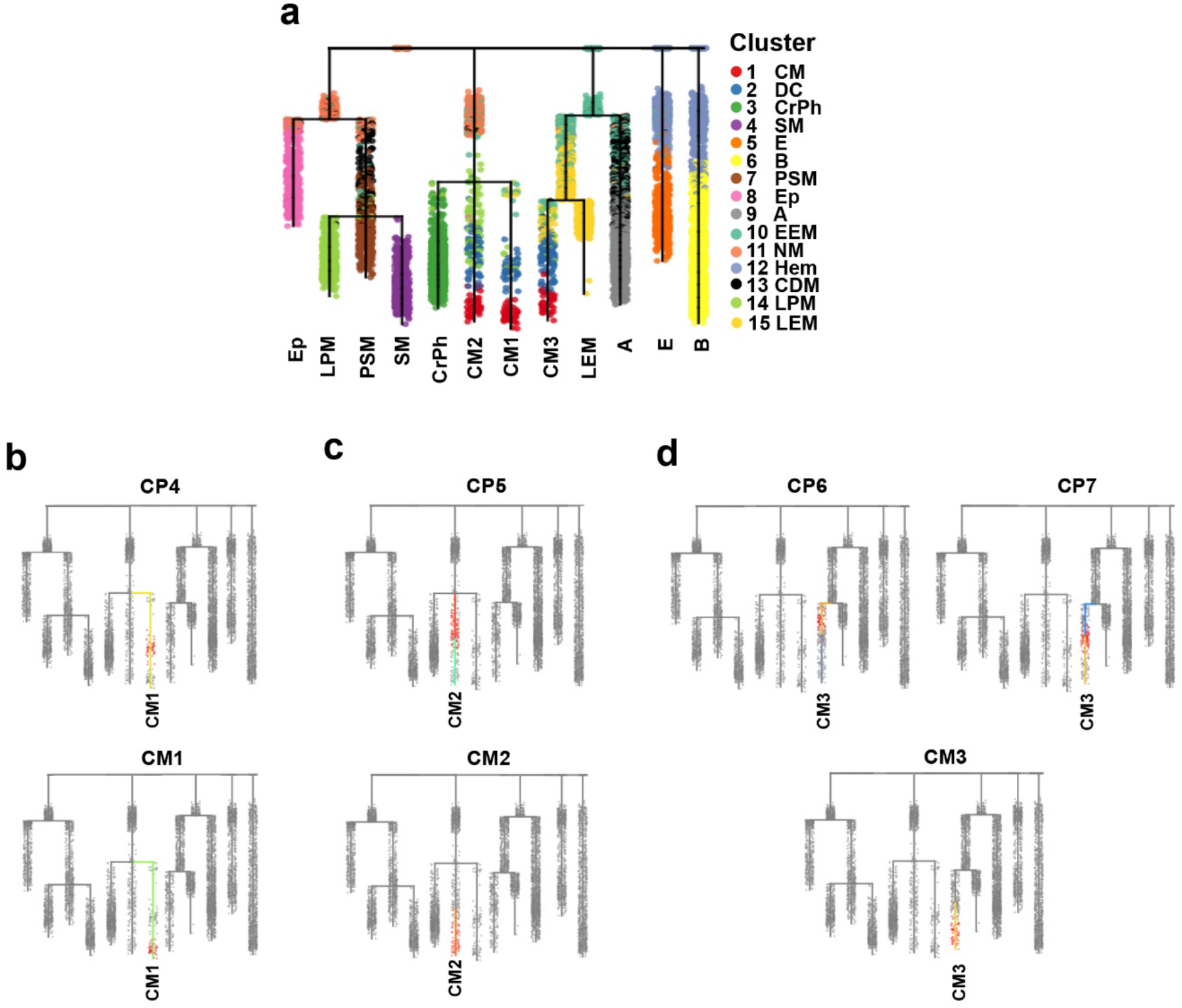
Cardiac subcluster cell types are located in specific branches of *Mesp1*-Cre URD developmental lineage tree. **a-d**, Each identified cardiac subcluster from Figure 2**c** is projected onto the (**a**) URD developmental lineage tree from Figure 3**a**. Projections of these subclusters show that they reside in specific regions of the (**b**) CM1, (**c**) CM2 and (**d**) CM3 branches.

**Extended Data Figure 6.**
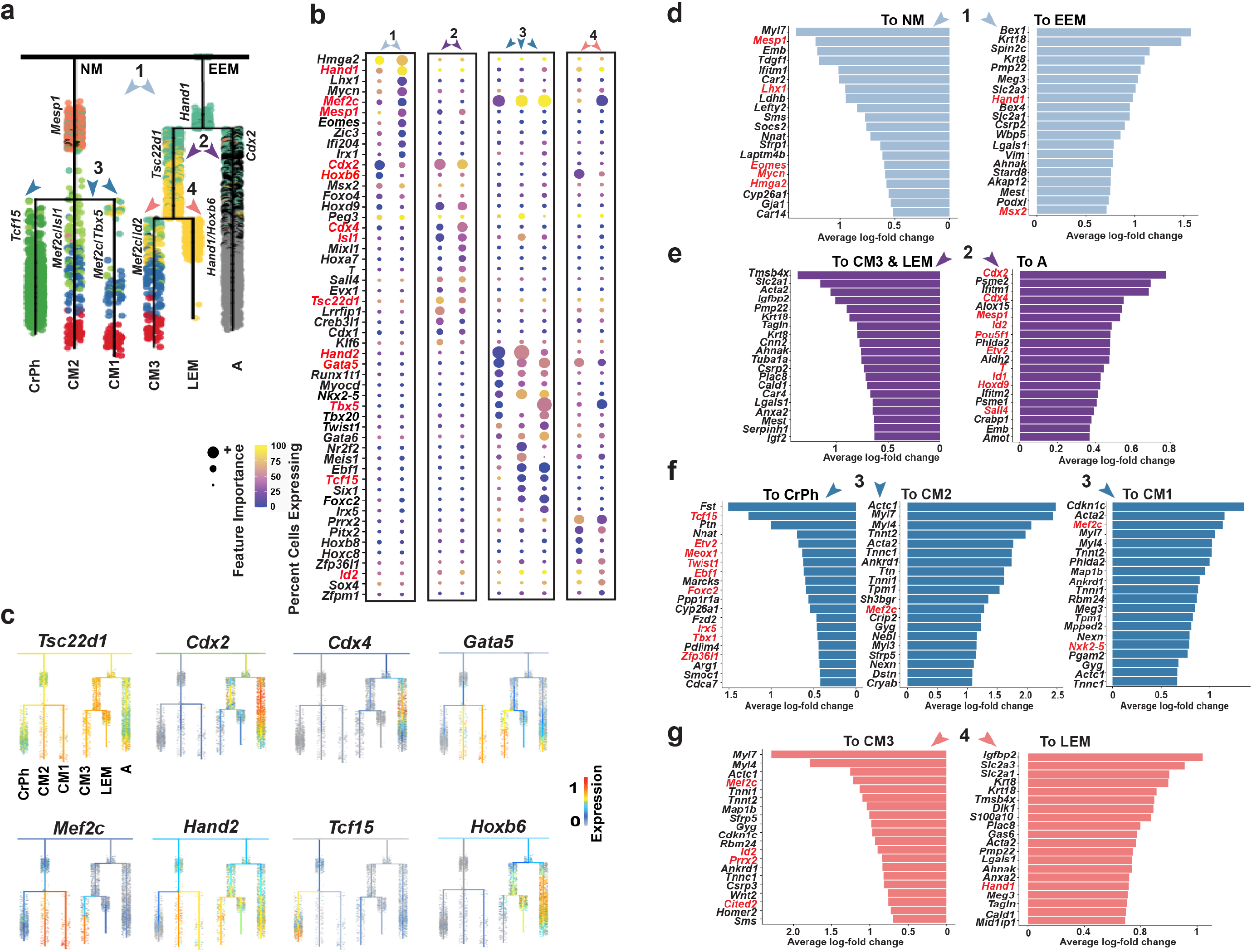
Branch point analyses of *Mesp1*-Cre cardiac lineage trajectories reveal molecular pathways for the development of distinct cardiac cell types. **a**, The cardiomyocyte-related branches of the URD developmental tree display the branch point decisions taken by cells differentiating into cardiomyocytes. Representative important transcription factors as determined in **b** are indicated on the corresponding branches of the URD developmental lineage tree. Cells are colored by their identity (see Figure 1) and numbers indicate each designated branch point analyzed. Arrowheads point to daughter branches. **b**, A random forest model was applied to predict the importance of individual transcription factors in directing cells to specific daughter branches at branch points labeled in **a**. The top ten transcription factors per branch, ranked by their importance, are shown as noted by size of dots, which are colored by the percentage of cells in each contrasting class/cells in the daughter branch just after the branch point. Red labeled genes indicate representative transcription factors for each branch. **c**, The expression of representative important transcription factors in **b** is projected onto the cardiomyocyte-related branches of the URD developmental lineage tree (also see Figure 3**c**). **d-g**, Bar plots display the most differentially expressed genes between the contrasting classes/cells in the daughter branch just after each respective branch point (as indicated by numbers and arrowheads). Transcription factors are labeled in red. Numbers and colored arrowheads indicate branch points marked in panel (**a**).

**Extended Data Figure 7.**
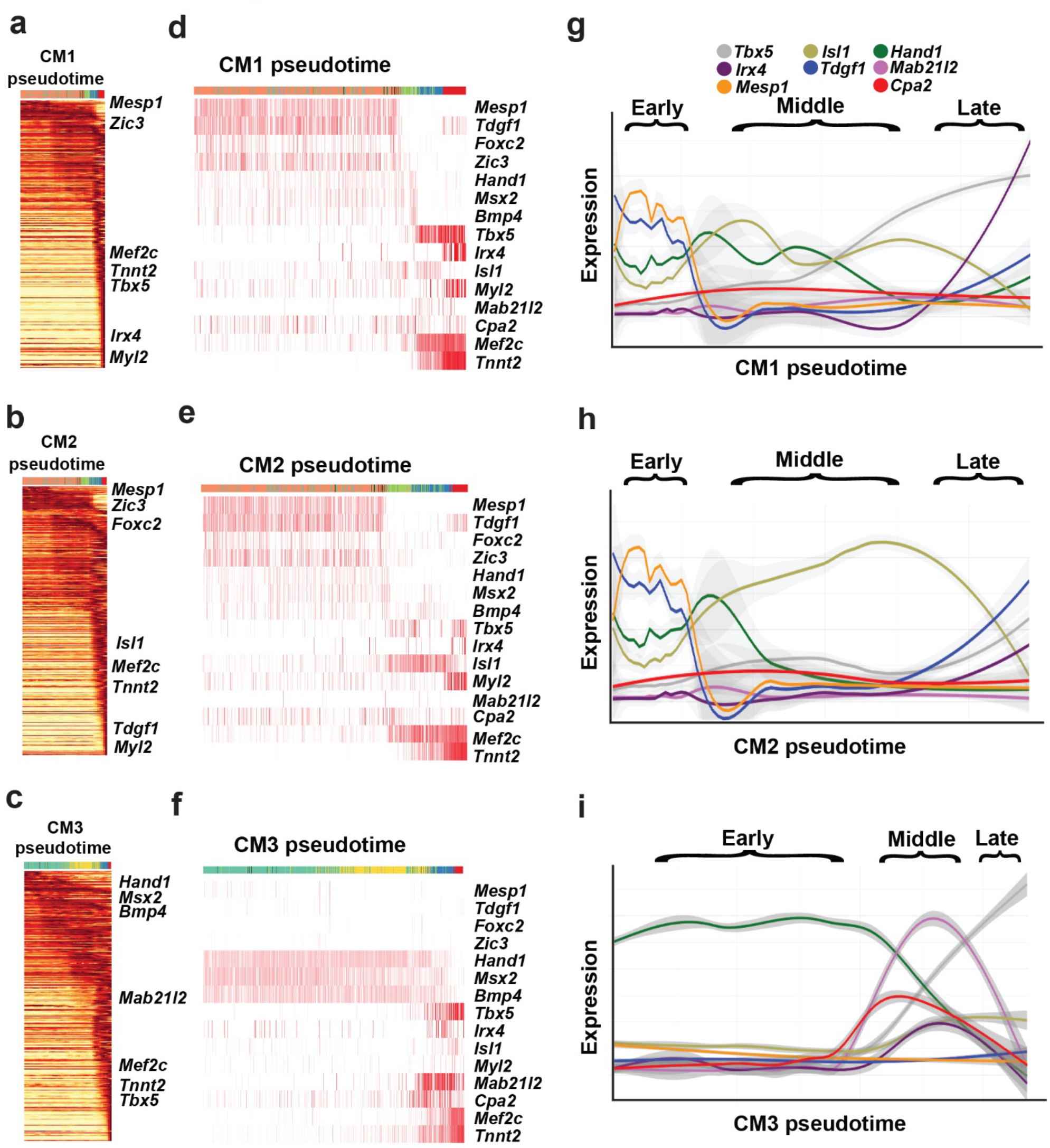
Pseudotime analyses of URD developmental lineage tree reveal the major developmental states and gene expression dynamics for each cardiomyocyte subcluster cell type. **a-c**, Heatmaps of differentially expressed genes for (**a**) CM1, (**b**) CM2 and (**c**) CM3 trajectories are displayed according to the pseudotime for each respective trajectory. **d-f**, Heatmaps of selected genes for (**d**) CM1, (**e**) CM2 and (**f**) CM3 trajectories are displayed according to the pseudotime for each respective trajectory. Corresponding cell types that are ordered by pseudotime and colored according to their cell identities in Figure 1 are shown at the top of each heatmap for **a-f**. **g-i**, Gene expression of key markers for each cardiomyocyte subcluster trajectory is plotted along the pseudotime for each of these trajectories (**g** – CM1, **h** – CM2, **i** – CM3). Colored lines indicate each gene examined (see legend above plots). Brackets signify pseudotime stages for each cardiomyocyte subcluster pseudotime plot.

**Extended Data Figure 8.**
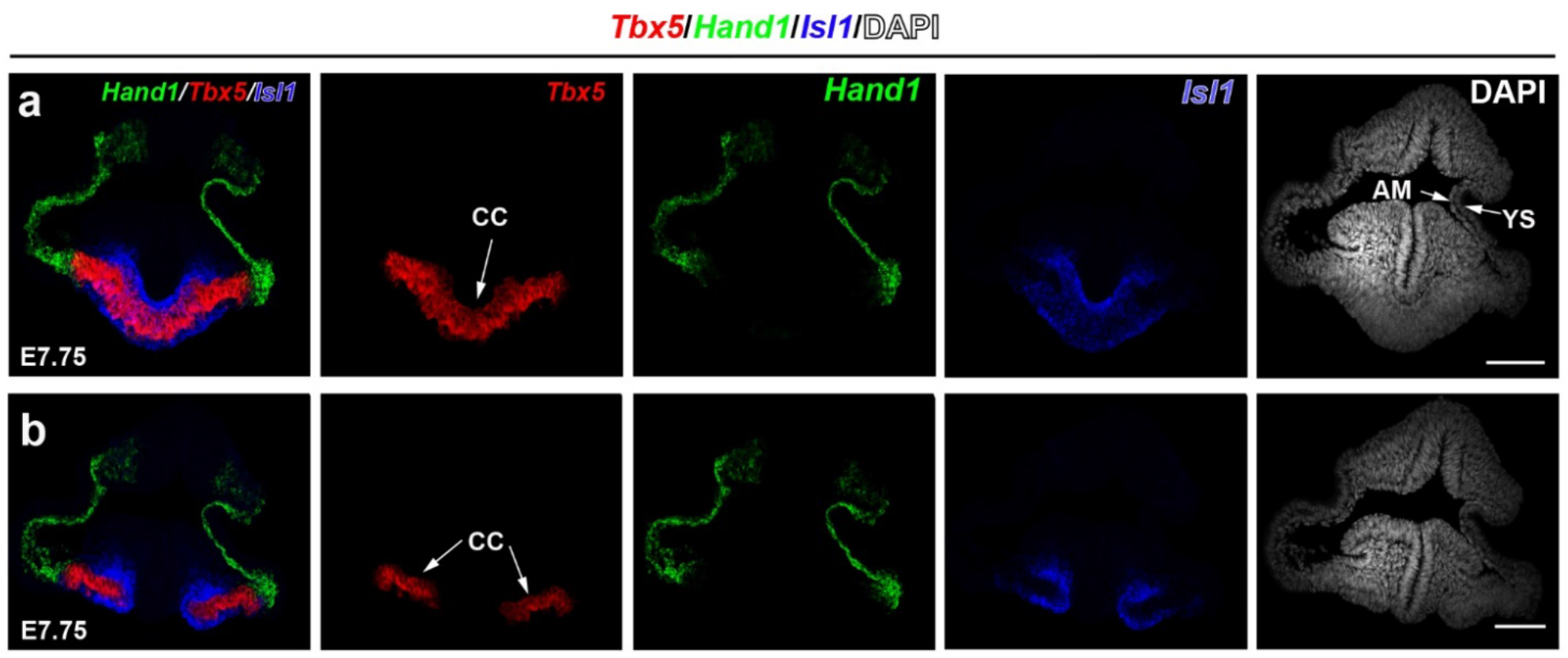
*Hand1, Tbx5* and *Isl1* are expressed in specific domains of the crescent region. **a**, **b**, RNAscope ISH studies on transverse sections of E7.75 embryos reveal that *Hand1, Tbx5* and *Isl1* are expressed in distinct but complementary domains within the crescent region. Furthermore, *Hand1* is expressed in additional areas that are outside but contiguous with the *Tbx5*-expressing cardiac crescent (CC). **a**, **b** panels show two different levels of the crescent region. n = 3 embryos. Scale bars, 100 μm. AM, Amnion; YS, Yolk Sac.

**Extended Data Fig. 9.**
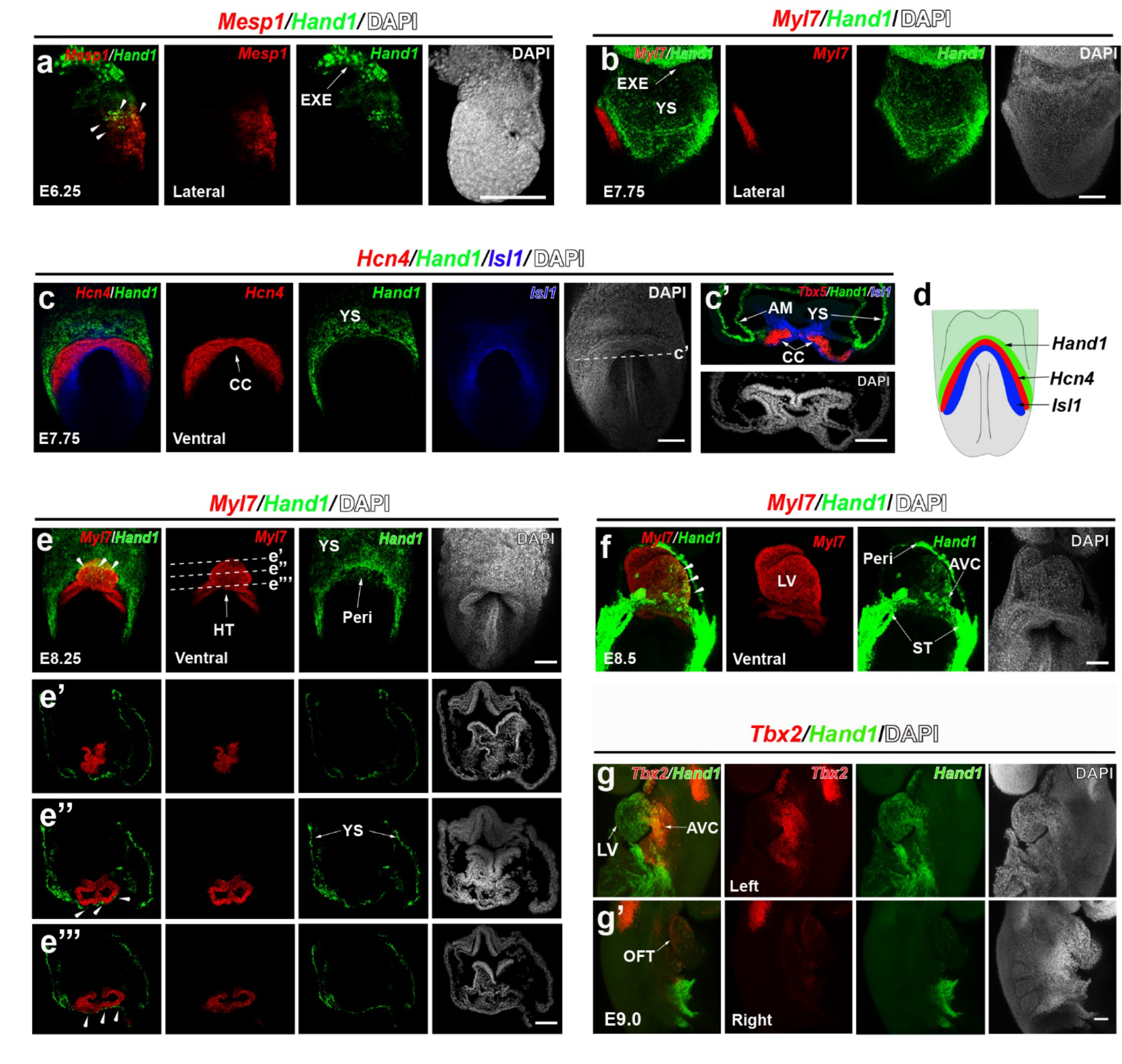
*Hand1* exhibits spatiotemporally dynamic expression in embryonic and extraembryonic tissues during embryogenesis. RNAscope *in situ* hybridization (ISH) studies were performed across different stages of mouse development to examine the dynamic expression of *Hand1* during embryogenesis. **a**, *Hand1* and *Mesp1* are co-expressed at the embryonic and extraembryonic boundary (arrowheads) in E6.25 embryos. **b**, *Hand1* is expressed in a region that is outside but contiguous with the *Myl7* expressing cardiac crescent in E7.75 embryos. **c**, **d**, *Hand1* expression complements *Hcn4* and *Isl1* expression in the crescent region at E7.75. **c’**, Inset shows transverse section of **c** at dashed line. **d**, Diagram illustrates *Hand1, Hcn4* and *Isl1* expression in the crescent region as shown in **c**. **e**, *Hand1* is expressed in the yolk sac and pericardium (arrowheads) which overlay the heart tube as detected by *Myl7* expression, but is not expressed in differentiated cardiomyocytes in E8.25 embryos. **e’**, **e’’**, **e’’’** Insets show transverse serial sections of the heart tube at corresponding dashed lines in **e**. **f,** *Hand1* is strongly expressed in the ST and pericardium, but weakly expressed in *Myl7*+ differentiated cardiomyocytes in the primitive LV and AVC (arrowheads) in E8.5 embryos. **g**, **g’**, *Hand1* is expressed in the AVC (*Tbx2+*) and LV, but is not expressed in the OFT in E9.0 embryos as shown in (**g**) left and (**g’**) right lateral views. n = 3 per panel. Scale bars, 100 μm. AM, Amnion; AVC, Atrioventricular Canal; CC, Cardiac Crescent; EXE, Extraembryonic Ectoderm; Heart tube, HT; OFT, Outflow Tract; Peri, Pericardium; ST, Septum Transversum; LV, Left Ventricle; YS, Yolk Sac.

**Extended Data Figure 10.**
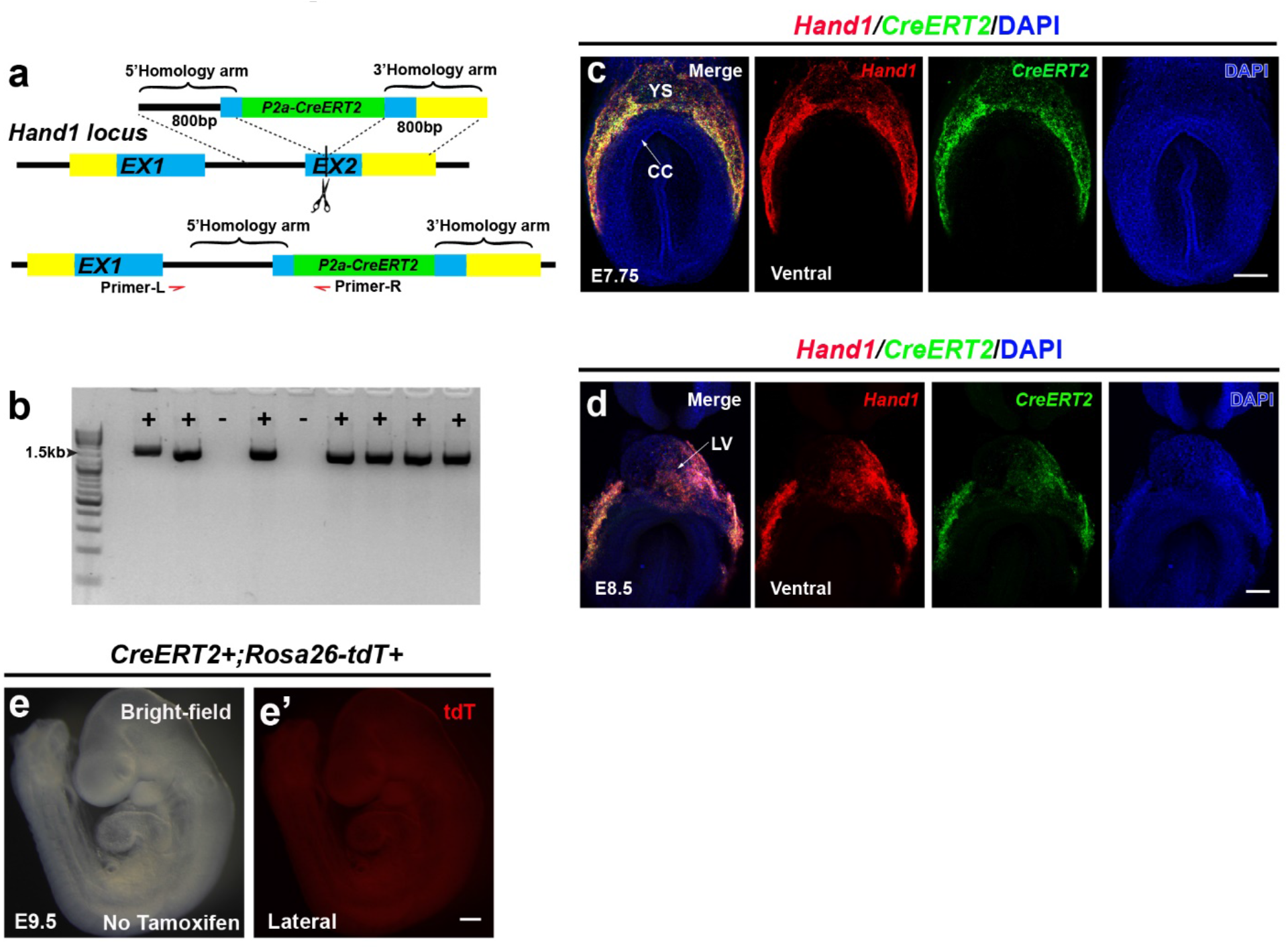
*Hand1-CreERT2* mouse line was generated by targeting a *P2a-CreERT2* construct into the second exon of the *Hand1* locus. **a**, Schematic illustrates the targeting strategy to create the *Hand1-CreERT2* mouse line. **b**, PCR analysis shows the genotyping for the identification of wildtype and *Hand1-CreERT2* alleles. **c**, **d**, RNAscope *in situ* hybridization studies reveal that the expression of *CreERT2* from *Hand1-CreERT2* mouse embryos recapitulates the endogenous expression of *Hand1* at (**c**) E7.75 and (**d**) E8.5. n = 3 per panel. Scale bars, 150 μm. **e**, In the absence of tamoxifen, no *Hand1-CreERT2* genetically-labeled tdT+ cells were observed in the E9.5 *Hand1-CreERT2; Rosa26-tdT* embryos. n = 5 embryos. Scale bars, 300 μm. CC, Cardiac Crescent; LV, Left Ventricle; YS, Yolk Sac.

**Extended Data Figure 11.**
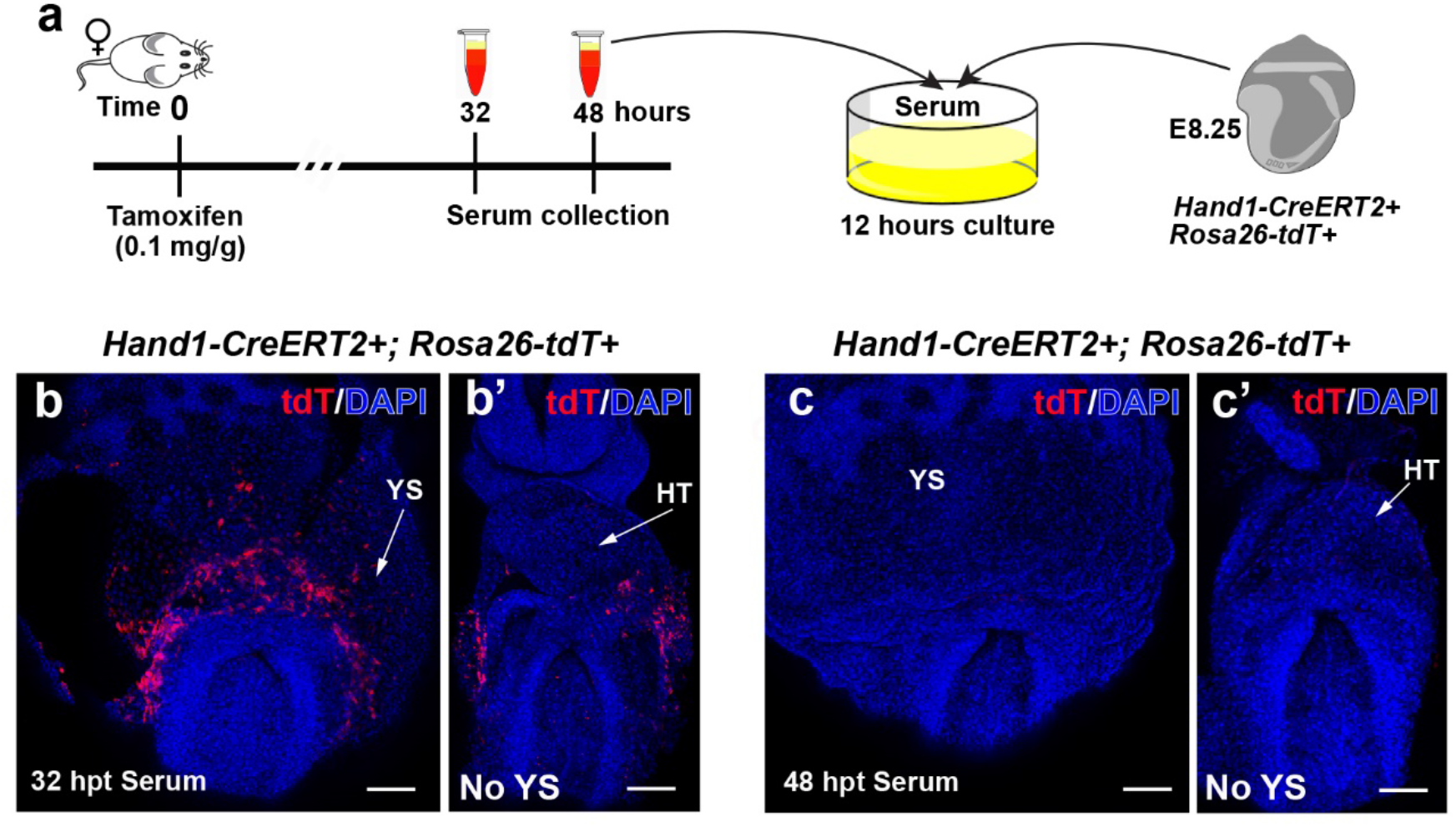
Tamoxifen can induce recombination up to 32 hours after treatment. **a**, Schematic illustrates experimental design for testing the perdurance of tamoxifen after treatment. Wild-type Black Swiss adult females were given tamoxifen at a 10.1 mg/g dose, and serum was collected at 32 and 48 hours post tamoxifen treatment (hpt). E8.25 *Hand1-CreERT2; Rosa26-tdT* embryos were cultured in collected sera for 12 hours. **b**, **b’**, These *Hand1-CreERT2; Rosa26-tdT* embryos cultured in serum collected 32 hours after tamoxifen treatment exhibited some *Hand1-CreERT2* genetically-labeled tdT+ cells in the yolk sac but not in the heart tube. n = 3. Scale bars, 150μm. **c**, **c’**, However, no genetically-labeled tdT+ cells were observed in *Hand1-CreERT2; Rosa26-tdTomato* embryos cultured in serum collected 48 hours after tamoxifen treatment. n = 4. Scale bars, 150μm. The extraembryonic tissue and part of the pericardium tissue were removed in **b’**, **c’** to show the underlying heart tube. HT, Heart Tube; YS, Yolk Sac.

**Extended Data Figure 12.**
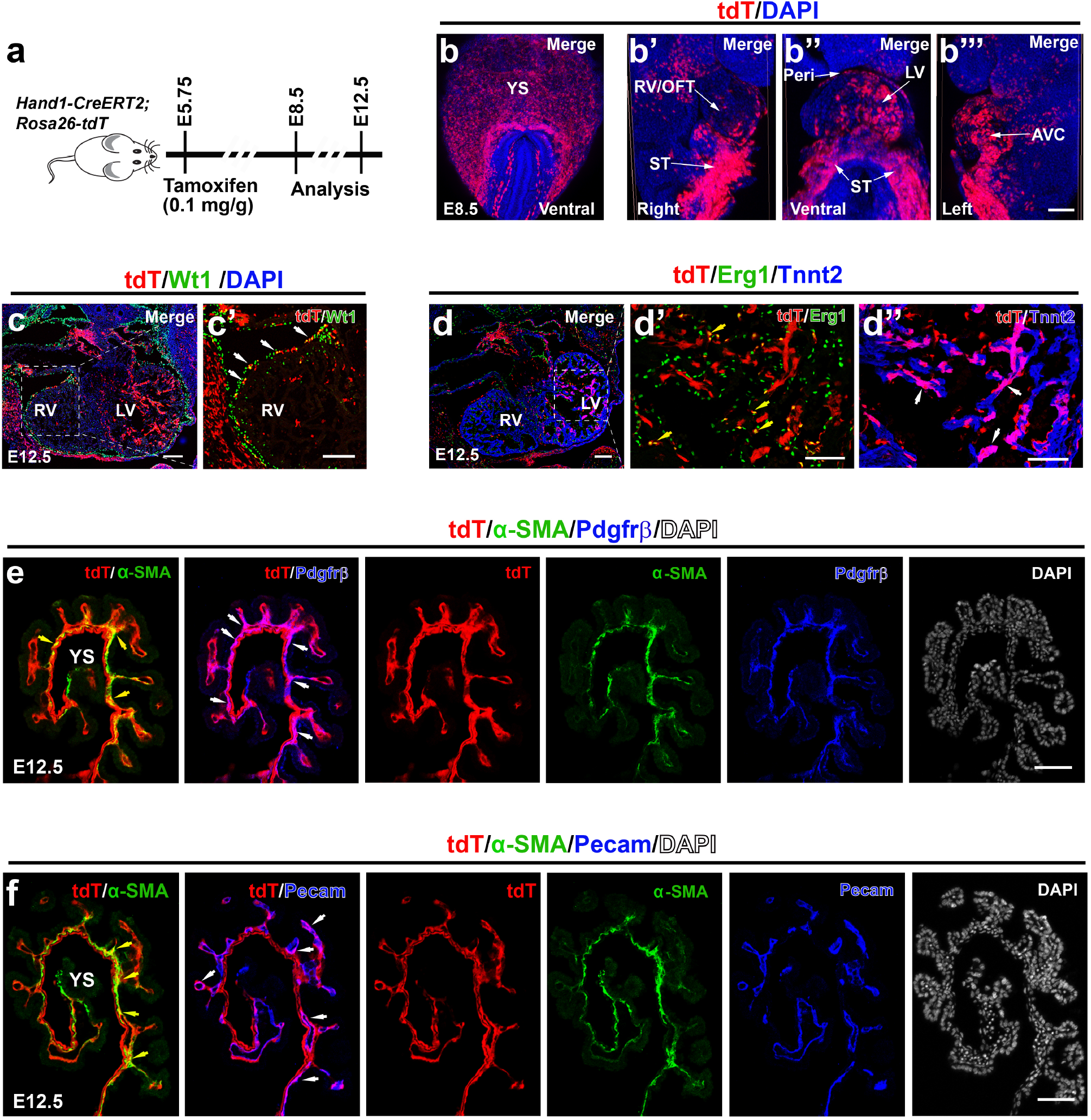
Lineage tracing studies show that early gastrulating *Hand1*+ cells contribute specifically to left ventricular cardiomyocytes, epicardial and few endocardial cells in the heart as well as endothelial cells, vascular support cells and mesothelial cells in the yolk sac. **a**, Schematic illustrates the experimental strategy that was used to assess the contribution of E5.75 genetically-labeled *Hand1-CreERT2; Rosa26-tdT* cells to E8.5 and E12.5 embryos. **b**, Whole mount embryo imaging shows that these *Hand1-CreERT2* genetically-labeled tdT*+* cells contribute to the yolk sac and developing heart at E8.5. **b’**, **b’’**, **b’’’**, The yolk sac (YS) and part of the pericardium tissue were removed in these panels to view the developing heart. **c-f**, Immunohistochemistry of cross-sectioned *Hand1-CreERT2; Rosa26-tdT* embryos at E12.5 reveals that *Hand1-CreERT2* genetically-labeled tdT+ cells contribute to epicardial (**c**, **c’**, Wt1, white arrowheads), myocardial (**d**, **d’’**, Tnnt2, white arrowheads) and few endocardial cells (**d**, **d’**, Erg1, yellow arrowheads) in the heart as well as (**e**, **f**) smooth muscle cells (α-SMA, yellow arrowheads), (**e**) mesothelial cells (Pdgfrβ, white arrowheads) and (**f**) endothelial cells (Pecam, white arrowheads) in the yolk sac. **c’**, **d’/d’’**, Insets are magnification of **c**, **d** boxed area, respectively. n = 3 per panel. Scale bars, 100 μm. AVC, Atrioventricular Canal; LV, Left Ventricle; OFT, Outflow Tract; Peri, Pericardium; ST, septum transversum; RV, Right Ventricle; YS, Yolk Sac.

**Extended Data Figure 13.**
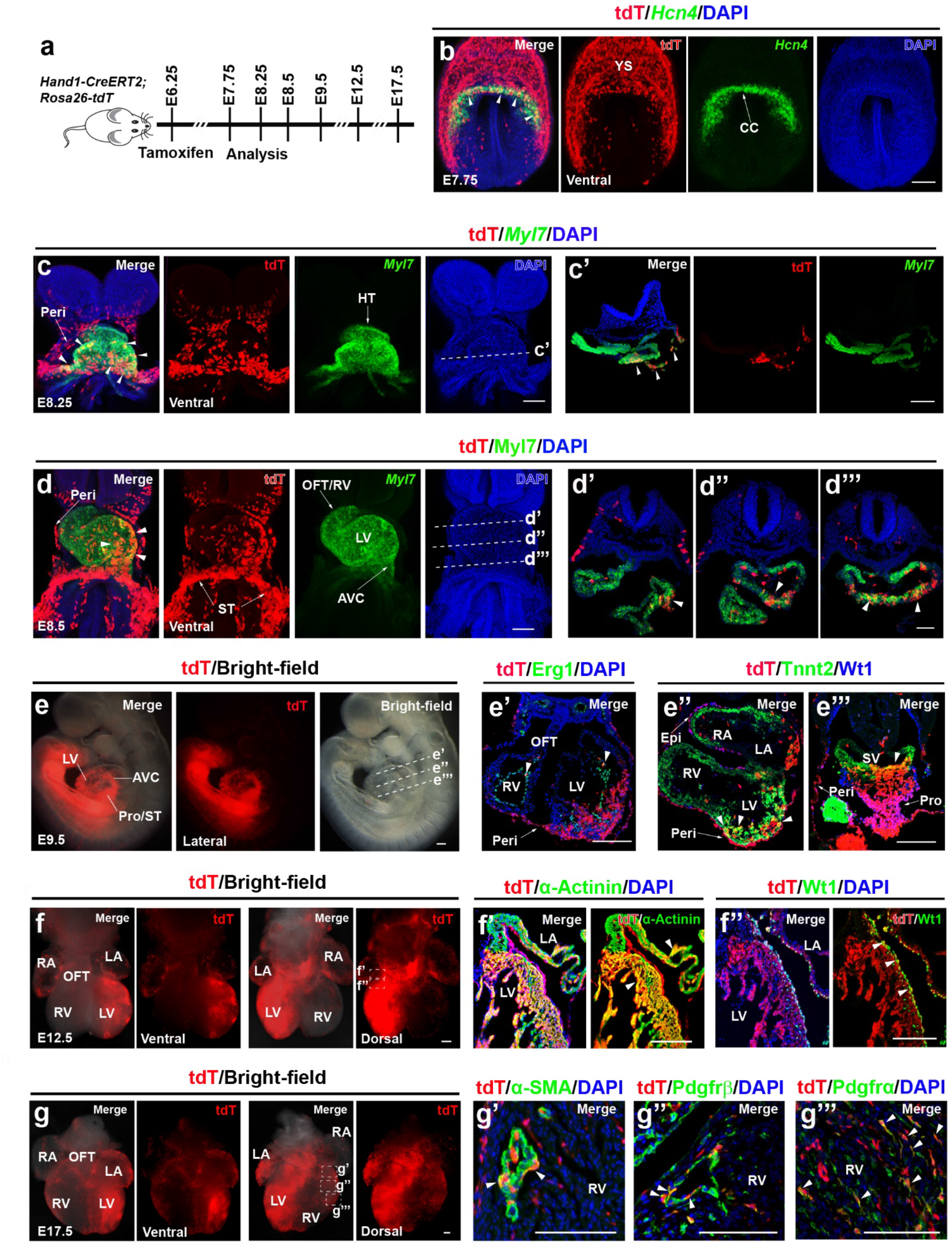
Lineage tracing studies marking *Hand1+* progenitors at E6.25 reveal that these cells contribute to first heart lineage cardiomyocytes and serosal mesothelial lineages (pericardial, epicardial cells) in the heart. **a**, Schematic outlines the experimental strategy for *Hand1-CreERT2* genetic fate mapping studies shown in **b-g**. Tamoxifen was given at E6.25, and *Hand1-CreERT2*; *Rosa26-tdT* embryos were examined for tdT localization at E7.75, E8.25, E8.5, E9.5, E12.5 and E17.5. RNAscope *in situ* hybridization and immunohistochemistry of whole mount and cross sections of these embryos (as indicated in each panel) reveal the contribution of *Hand1-CreERT2* genetically-labeled tdT+ cells at (**b**) E7.75, (**c**) E8.25, (**d**) E8.5 (**e**) E9.5, (**f**) E12.5 and (**g**) E17.5. **c’**, **d’-d’’’**, **e’-e’’’**, Insets show transverse sections of **c**, **d**, **e** at indicated dashed lines, respectively. **f’**, **f’’** and **g’-g’’’**, Inset shows representative coronal sections of **f** and **g** at indicated dashed boxes, respectively. Arrowheads point to tdT+ cells expressing (**b**) *Hcn4*, (**c**, **d**) *Myl*7, (**e’**) Erg1, (**e’’**) Tnnt2, (**e’’’**, **f’’**) Wt1, (**f’**) α-Actinin, (**g’**) α-SMA, (**g’’**) Pdgfrβ and (**g’’’**) Pdgfrα. n = 3 for each condition. Scale bars, 100 μm. Embryos analyzed at E17.5 were given 0.05 mg/g tamoxifen, half the dose given to embryos analyzed at earlier timepoints. AVC, Atrioventricular Canal; CC, Cardiac crescent; Epi, Epicardium; HT, Heart tube; LA, Left Atrium; LV, Left Ventricle; RA, Right Atrium; RV, Right Ventricle; OFT, Outflow Tract; Peri, Pericardium; Pro, Proepicardium; ST, Septum transversum; SV, Sinus Venosus; YS, Yolk sac.

**Extended Data Figure 14.**
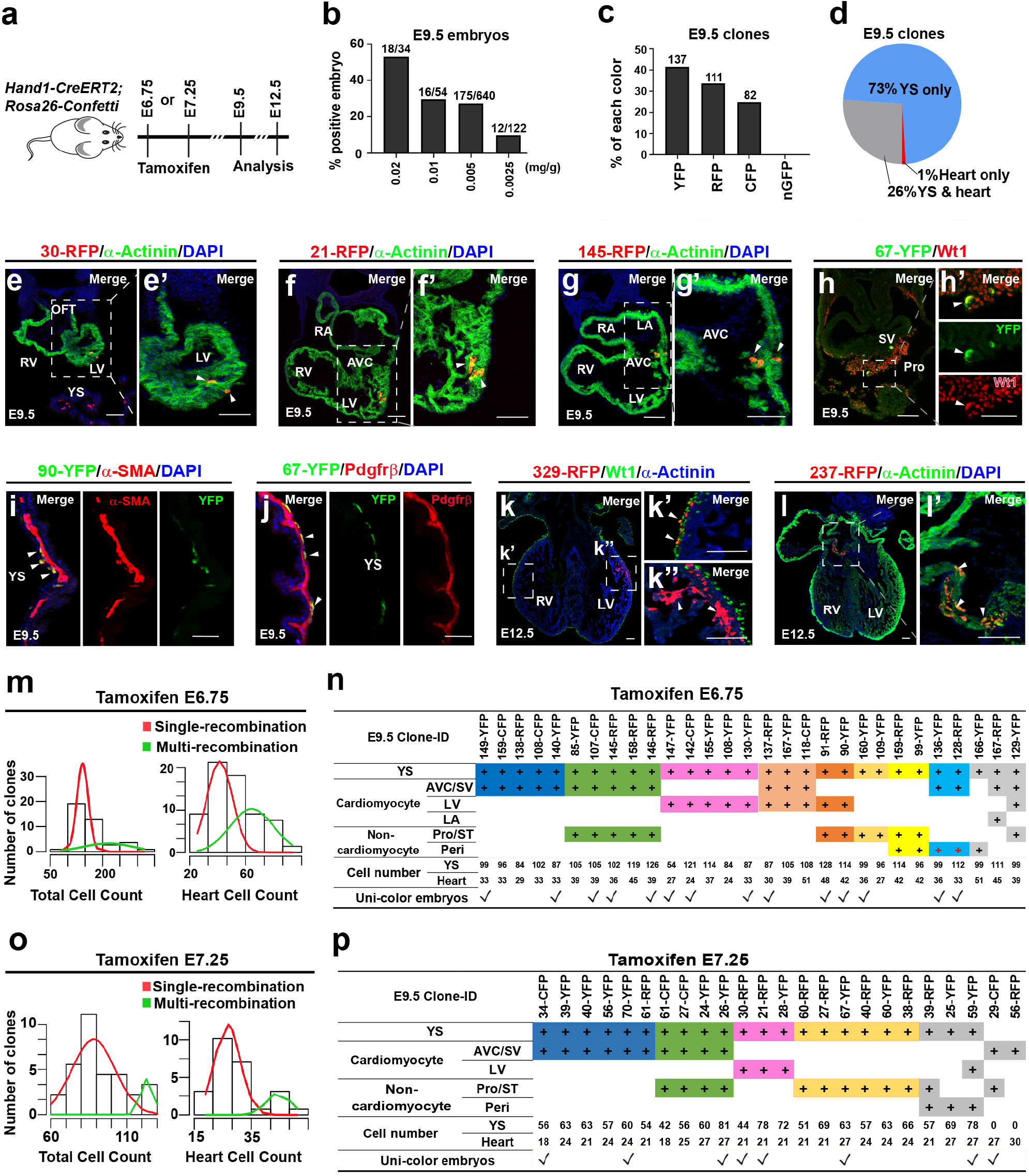
Quantitative analysis of clones and their cell type identification reveal the contribution of *Hand1+* progenitor clones to specific cell lineages of the heart and yolk sac. **a**, Schematic outlines experimental strategy for *Hand1-CreERT2; Rosa26-Confetti* clonal analyses in **b-p**. **b**, Bar graph reveals the percentage of E9.5 embryos that displayed fluorescence at titrated doses of tamoxifen. **c**, Bar graph displays the frequency of each fluorophore expressed in E9.5 *Hand1-CreERT2; Rosa26-Confetti* embryos that were induced with 0.005 mg/g tamoxifen. **d**, Pie chart shows the contribution of *Hand1-CreERT2; Rosa26-Confetti* clones to respective tissues. **e-l**, Immunohistochemistry of (**e-j**) E9.5 and (**k-l**) E12.5 *Hand1-CreERT2; Rosa26-Confetti* embryos reveals the contribution of *Hand1-CreERT2* genetically-labeled clones to specific cell types including (**e**, **f**, **g**, **k**, **l**) cardiomyocytes (α-Actinin) and (**h**, **k**) epicardial cells (Wt1) in the heart as well as (**i**) smooth muscle cells (α-SMA) and (**j**) mesothelial cells (Pdgfrβ) in the yolk sac. **e’-h’**, **k’**, **k’’**, **l’**, Insets are magnification of **e-h**, **k**, **l** boxed area. Arrowheads point to *Hand1-CreERT2; Rosa26-Confetti* labeled clones expressing (**e’**, **f’**, **g’**, **k’’**, **l’**) α-Actinin, (**h’**, **k’**) Wt1, (**i**) α-SMA, and (**j**) Pdgfrβ. ID number for each clone analyzed is indicated in panels. Scale bars = 100 μm. **m**, **o**, Histograms show the number of (**m**) E6.75 and (**o**) E7.25 genetically-labeled clones with a specific cell count (total or only in the heart). Gaussian distributions representing single (red line) and multiple (green line) recombinant events were modeled on this data. Based on these distributions, only clones that likely derived from a single recombination events were analyzed. These clones and their contributions to specific cell types are shown in (**n**) and (**p**), respectively. Unicolor embryos which are shown in Figure 5 are denoted by check mark on bottom of tables. AVC, Atrioventricular Canal; LA, Left Atrium; LV, Left Ventricle; RA, Right Atrium; RV, Right Ventricle; OFT, Outflow Tract; Peri, Pericardium; Pro, Proepicardium; ST, Septum transversum; SV, Sinus Venosus; YS, Yolk sac.

**Supplementary Table 1. Differential gene expression analyses assign cell identities to scRNA-seq *Mesp1*-Cre clusters.** Table displays differentially expressed marker genes for each cluster appearing in Figure 1 as defined in Methods. All tests are Wilcoxon rank sum tests. Column titles indicate the following information: gene – gene marker analyzed; cluster name – cluster expressing gene marker; p_val – unadjusted p-value of association with the cluster in contrast to all cells not in the cluster; avg_logFC average – average log2 change between the expression of the marker in the cluster versus all cells not in the cluster; pct.1 – percentage of cells in the marked cluster that express the gene at a non-zero level; pct.2 – percentage of cells not in the marked cluster that express the gene at a non-zero level; p_val_adj – multiple hypothesis adjusted p-value of association with the cluster in contrast to all cells not in the cluster; and cluster – numeric ID of the cluster as assigned in Figure 1.

**Supplementary Table 2. Differential gene expression analyses assign cell identities to cardiac subclusters.** Table displays differentially expressed genes for each cardiac subcluster appearing in Figure 2. All tests are Wilcoxon rank sum tests. Column titles indicate the following information: gene – gene marker analyzed; cluster name – cardiac subcluster expressing gene marker; p_val – unadjusted p-value of association with the subcluster in contrast to all cells not in the subcluster; avg_logFC average – average log2 change between the expression of the marker in the subcluster versus all cells not in the subcluster; pct.1 – percentage of cells in the marked subcluster that express the gene at a non-zero level; pct.2 – percentage of cells not in the marked subcluster that express the gene at a non-zero level; p_val_adj – multiple hypothesis adjusted p-value of association with the subcluster in contrast to all cells not in the subcluster.

**Supplementary Table 3. Gene expression analyses reveals genes differentially expressed in branches at each branchpoint analyzed in the cardiomyocyte trajectories.** Tables display genes that are differentially expressed between branches at each branch point analyzed in Extended Data Figure 6. All tests are Wilcoxon rank sum tests. Each sheet shows genes that are differentially expressed at corresponding branch points as indicated in the sheet name. Column titles indicate the following information: gene – gene marker analyzed; p_val – unadjusted p-value of association between two branches analyzed at respective branch point; avg_logFC average – average log2 change of gene expression between the two indicated branches analyzed; pct.1 and pct. 2 – percentage of cells expressing the gene in each branch as indicated in the column header; p_val_adj – multiple hypothesis adjusted p-value of differential expression between the two branches.

**Supplementary Table 4. Primer and sequence for making or genotyping *Hand1-CreERT2, Rosa26-tdTomato* and *Rosa26-Confetti* mice are shown.** PCR primer, *CreERT2* and gRNA sequences are provided for the making or genotyping of *Hand1-CreERT2, Rosa26-tdT* and *Rosa26-Confetti* mice.

## Methods

### Animal models

Animal studies were conducted in strict compliance with protocols approved by the Institutional Animal Care and Use Committee of the University of California, San Diego (UCSD) (A3033-01) and *the Guide for the Care and Use of Laboratory Animals* published by the National Institutes of Health. Mice were kept in IVC disposable cages (Innovive), under a 12-hour light cycle and bred on the Black Swiss background (Charles River Labs). We used *Mesp1-Cre, Rosa26-tdT* and *Rosa26-Confetti* mouse lines for our studies, which have been previously described^19,20,63^. The *Hand1-CreERT2* knock-in line was made as described^76^. Briefly, this procedure entailed using Gibson cloning to create a donor DNA fragment which contains a *P2a-CreERT2* sequence surrounded by 1600 bps of homology sequence to the second exon of *Hand1* (See Supplementary Table 4 for primer sequences). This fragment was fully sequenced in order to ensure that mutations had not been introduced. The donor DNA (0.6μM) with Cas9 protein (NEB #M0646T), crRNA and tracrRNA were injected in a 1:1:1:1 molar ratio into mouse zygotes by the UCSD Transgenic and Knockout Mouse Core (See Supplementary Table 4 for crRNA sequence). Four independent founders were recovered, of which three displayed strong Cre activity and expression. No differences were detected among these three founders, which were further propagated and used for experiments. Additionally, RNAscope *in situ* hybridization (ISH) confirmed that expression of *CreERT2* from these mice recapitulates expression of endogenous *Hand1* (Fig. 4**e**, Extended Data Fig. 10**c, d**). For genotyping, genomic DNA was extracted by adding 75 µl of 25 mM NaOH, 0.2 mM EDTA to a 2 mm tail clipping and heating at 98°C for 30 minutes. The solution was then neutralized by adding 75 µl of 40 mM Tris-HCl (pH 5.5). A 1:50 dilution of genomic DNA template was used for genotyping PCR. Primers matching sequences upstream of the left homology arm and in the *Cre* gene were used for genotyping (See Supplementary Table 4 for primer sequences.).

### Embryo dissection and scRNA-seq library generation

To prepare single cells for scRNA-seq, *Mesp1-Cre; Rosa26-tdT* genetically-labeled embryos at E7.25, E7.5, E7.75 and E8.25 were dissected in cold sterile 1 X PBS without Ca^2+^, Mg^2+^ under a stereo microscope. Embryos were staged based on their morphology^77^. The Reichert’s membrane and ectoplacental cone were removed, and *Mesp1-Cre; Rosa26-tdT* genetically-labeled embryos were selected and imaged. The yolk sac was removed from two of the three E8.25 embryos that were processed in order to enrich for cardiac cells. Individual embryos were placed into a 1.5 ml microfuge tube and incubated in 0.25% Trypsin-EDTA (Gibco, Catalog # 25200056) at 37°C with inversion every two minutes for 30 min until no visible tissue remained. The solution was pipetted once with a p1000 and neutralized by adding 0.75 ml DMEM containing 10% FBS (Gibco). Cells were then passed through a 100-μm cell strainer (BD Biosciences, Catalog # 352360) and single tdT+ cells were obtained by fluorescence-activated cell sorting (FACS) on a BD Influx Cell sorter (BD Biosciences). Living cells were gated on FSC, SSC, DAPI- and tdT+. After sorting, cells were centrifuged at 300g for 4 minutes and pooled or kept as individual embryos. Libraries were prepared using the Chromium Single Cell 3′ Library and Gel Bead Kit v2 (PN-120237) and Chromium i7 Multiplex Kit (PN-120262) according to instructions from 10X Genomics (https://www.10xgenomics.com/resources/user-guides/). Prior to sequencing cDNA, libraries were verified by the D1000 ScreenTape system (Agilent) and quantified via Qubit™ Flex Fluorometer (Thermofisher, Catalog # Q33327). All libraries were sequenced twice on the HiSeq 4000 (Illumina) at the UCSD genomics core. An initial shallow sequencing run was done for quality control and to determine the number of cells captured. An individual sample was excluded from further analysis due to a low number of reads per cell (60%) as analyzed by Cell Ranger (10X genomics). A second deeper sequencing run was subsequently performed ensuring an approximate equal read depth per cell across the samples, resulting in an average of 60,450 UMIs (unique molecular identifiers) per cell and an average sequence saturation of 65.7% (Extended Data Fig. 1**b**).

### Data processing and clustering

Reads were analyzed with the Seurat library (version 3.1.5). The data was read into the R (version 3.5.3) computing environment and log normalized using the Seurat library’s NormalizeData() function with default parameters. Cells with more than 5 percent mitochondrial gene reads or less than 25,000 UMI were excluded. This analysis excluded approximately 1,400 cells (Extended Data Fig 1**a, b**). We then calculated the principal components of the data and used the first 10 principal components to calculate tSNE projections. Individual samples visualized in these tSNE projections revealed that samples overlapped, thus indicating a lack of batch effect (Extended Data Fig. 1**c**). In order to discover an optimal number of clusters for analysis, we also used ten principle components and calculated k-means clustering for k = 8 to 25. For each clustering solution k, we observed the average silhouette score. We observed local maxima at k = 10, 12 and 15, and chose k = 15 for subsequent analysis (Extended Data Fig. 1**d**). For each of these clusters, we identified genes that were expressed at higher levels in that cluster compared to all other cells using default parameters for Seurat’s FindMarkers function. The complete list of these genes along with the clusters that they represent can be found in Supplementary Table 1. These genes were examined more closely in order to assign a cell identity to each cluster. The most informative markers appear in Figure 1**f**.

### Lineage inference

In order to infer the developmental relationships between cells in our study, we employed the R package URD (version 1.1.0)^28^, which requires the user to declare certain cells to be part of the root or the tips of the cell lineage tree. URD then traces routes through a cell-cell nearest neighbor graph from the tip cells back to the root cells producing a tree-topology that summarizes the consensus routes from each tip back to the root cells. We used all cells from our earliest stage E7.25 (No bud) as the root of the tree, and cells of clusters that contained the most differentiated cell-types at the latest stage E8.25 (1-4 somite) as the tips. For the URD in Figure 2**a**, cells from E8.25 embryos from the following clusters were defined as tips: Allantois, A; Blood, B; Cardiomyocyte, CM; Cranial pharyngeal mesoderm, CrPh; Endothelium, E; Epithelium, EP; Lateral plate mesoderm, LPM; Late extraembryonic mesoderm, LEM; Pre-somitic mesoderm, PSM and Somite mesoderm, SM. For the URD in Figure 3**a**, the same clusters were defined as tips except the CM tip was split into three tips, based on the sub-clusters (CM1, CM2, CM3) defined by re-clustering only the cardiomyocyte branch as described in the results.

To compare the Pijuan-Sala et al. data^27^ with our own, we first limited the analysis of their data to the developmental stages that we analyzed (E7.25 - E8.5). Analogous clusters between our data and theirs were determined by identifying the clusters which had the greatest number of identical marker genes. Marker genes for clusters in Pijuan-Sala et al. were defined as genes significantly associated to each cluster (adjusted p-val <0.05) as reported on their data portal (https://marionilab.cruk.cam.ac.uk/MouseGastrulation2018/). Marker genes in our data were determined as described above in Data processing and clustering section. To identify the clusters with the greatest number of identical marker genes between our dataset and the Pijuan-Sala et al. dataset, the number of marker genes from each of our clusters that match each of their clusters was divided by the total number of marker genes for the relevant cluster in our dataset. This value represents the size-normalized overlap between clusters in the two datasets and was plotted as a heatmap (Extended Data Fig. 3**c**). Clusters with maximal overlap were considered analogous between the two datasets. An URD tree was then created using analogous root (all E7.25 cells) and tip clusters as defined above. The cardiomyocyte branch of this URD tree was shown (Extended Data Fig. 3**d**).

### Branch point differential analysis

At each branch point along the three cardiomyocyte developmental trajectories (Figure 3**a**, CM1, CM2, CM3), we used a Random Forest model^44^ to identify transcription factors likely to be responsible for cells choosing one branch over another. For this algorithm, we defined contrasting classes of cells as the first ∼300 daughter cells for each branch that was compared at corresponding branch points. The feature set was defined as transcription factors (as identified by the Gene Ontology, DNA Binding, GO:0003677 term and manual annotation^78^). We used these contrasting classes and the feature set in the R library randomForest’s main function randomForest() using default parameters (except the importance=TRUE option was set to return the feature importance measures). The importance measure used here is the mean decrease in accuracy measure (the default for RandomForest()), which quantifies the decrease in prediction accuracy of a class when the variable in question is randomly permuted. The top ten most important transcription factors that determined each class were plotted (Extended Data Fig. 6**b**). The differential analysis between branches at each branch point used the contrasting classes of cells defined above, but examined all genes (instead of just transcription factors) using Seurat’s FindMarkers() function, with default parameters. The top twenty differentially positive expressed genes for each class as determined by their log-fold change were plotted (Extended Data Fig. 6**d-g**).

### Pseudotime Trace Analysis

We used URD-defined pseudotime^28^ for our pseudotime analysis: which is the average number of transitions over edges of the nearest neighbor graph required to reach each cell from the root. In order to produce the pseudotime traces, we ordered cells along each lineage according to the URD-inferred pseudotime using URD’s geneCascadeProcess() function (Extended Data Fig. 7**a-c**). The scaled expression from Seurat of each marker gene in each cell was plotted as a heatmap (Extended Data Fig. 7**d-f**) and as a smoothed spline (Extended Data Fig. 7**g-i**). Pseudotime stages (early, middle, late) were defined based on gene expression peak coherence in the smooth spline plots.

### Tamoxifen treatment

To determine the developmental stage of embryonic development during which tamoxifen treatment was administered, noon on the day of the vaginal plug was assumed to be E0.5. For lineage tracing studies, tamoxifen (Sigma, T5648-1G, 0.1 mg/g body weight) was fed to pregnant mice by gavage, except for embryos harvested at E17.5 when a lower dose of tamoxifen was used (0.05 mg/g body weight). For the clonal analysis, tamoxifen was administered by intraperitoneal injection.

### Lineage tracing and clonal analysis

For lineage and clonal analyses, *Hand1-CreERT2* mice were crossed with *Rosa26-tdT* or *Rosa26-Confetti* mice respectively. Genetically-labeled embryos were identified using a fluorescent stereo microscope (ZEISS AXIO Zoom.V16 or LEICA M205 FA). Embryos younger than E9.5 were imaged using a confocal microscope (Nikon C2), while embryos older than E9.5 were imaged with a fluorescent stereo microscope and then imaged with the confocal microscope after sectioning. Embryos were embedded and sectioned into 10 or 20 μm sections for lineage tracing or clonal analysis, respectively. To determine the number of cells in a clone, sections from an individual embryo were processed and distributed evenly across three slides for E9.5 embryos, or five slides for E12.5 hearts. All cells from an individual clone on one slide were then counted. The total number of cells per clone was then calculated by multiplying the number of cells in a clone on a single slide by the number of slides.

Tamoxifen perdurance was determined by incubating E8.25 embryos in serum collected from Black Swiss females 32 or 48 hours after they were given tamoxifen (0.1 mg/g). Serum was collected by centrifuging (2x 400g for 6 mins) blood collected via retro-orbital bleeding and then frozen at −80°C. Separately, E8.25 embryos were obtained from *Hand1-CreERT2* x *Rosa26-tdT* crosses without tamoxifen administration and dissected in 5% FBS/Fluorobrite DMEM media (ThermoFisher, Cat. no. A1896701 and 10082139) on a 37°C heated stage (Tokai Hit, TPi-SZX2AX). Care was taken to remove the Reichert’s membrane, but not the ectoplacental cone. Embryos were then incubated at 37°C in 5% CO_2_ for 12 hours in 2 ml of pre-warmed serum collected from tamoxifen-injected females. After incubation, embryos were fixed in 4% PFA and processed for immunofluorescence with anti-tdT antibody. Embryos which did not display tdT were genotyped to confirm that they contained both *Hand1-CreERT2* and *Rosa26-tdT* DNA. Three independent experiments were performed.

### RNAscope Fluorescent *in situ* hybridization

Whole-mount RNAscope fluorescent *in situ* hybridizations (ISH) were conducted using the RNA-scope Multiplex Fluorescent Reagent Kit v.2 (Advanced Cell Diagnostics, 323100) with several adaptations. Embryos were dissected in RNase-free 1X PBS and fixed in 4% PFA overnight at 4°C. Embryos were then washed 3 times in 0.1%Tween20/PBS (PBT), followed by dehydration into and then re-hydration from methanol using 5 minute 25%, 50%, 75% and 100% Methanol/PBT washes. Probe hybridization was performed at 50°C. For samples that were co-stained with antibodies after the RNAscope ISH, samples were incubated in 10% heat-inactivated donkey serum for 2 hours at room temperature prior to addition of primary antibody (see below) overnight at 4°C. The embryos were then washed 3x in PBT and incubated in secondary antibody (see below) for 2 hours in 4°C. Whole-mount embryos were imaged after mounting in 1% low melting point agarose in 35 mm glass bottom petri dishes (MatTek) using a confocal microscope (Nikon C2). After imaging, embryos were embedded and sectioned for further analysis as described in the Immunofluorescence, sectioning and image processing section. Catalog numbers for RNA-scope probes (ACDbio) used in this study: Cre-O4-C1, Cat No. 546951; Mm-Hcn4-C1, Cat No. 421271; Mm-Hand1-C1, Cat No. 429651; Mm-Hand1-C2, Cat No. 429651-C2; Mm-Isl1-C2, Cat No. 451931-C2; Mm-Mab21l2-C1, Cat No. 456901; Mm-Myl7-C3, Cat No. 584271-C3; Mm-Mesp1-C3, Cat No. 436281-C3; Mm-Nkx2-5, Cat No. 428241; Mm-Nkx2-5-C2, Cat No. 428241-C2; Mm-Nr2f2, Cat No. 480301; Mm-Sfrp5-C1, Cat No. 405001; Mm-Smoc2-C1, Cat No. 318541; Mm-Tbx5-C1, Cat No. 519581; Mm-Irx4-C1, Cat No. 504831; Mm-Tbx5-C2, Cat No. 519581-C2; Mm-Tdgf1-C1, Cat No. 506411.

### Immunofluorescence, sectioning and image processing

Immunofluorescence studies were conducted on cryosections of mouse embryos. Embryos were cryoprotected, mounted, sectioned and stained as we previously described^12^. The following primary antibodies were used: mouse anti-TNNT2 antibody (Invitrogen, Catalog # MA5-12960, 1:50), Chicken anti-GFP antibody (Abcam, ab13970, 1:300), Rabbit anti-ERG1 antibody (Abcam, ab92513, 1:300), Rabbit anti-WT1(Abcam, ab89901, 1:200), Rabbit anti-PDGFRα antibody (Abcam, ab203491, 1:200), Rabbit anti-PDGFRβ antibody (Abcam, ab32570, 1:200), Rat Anti-Mouse CD31 (BD Pharmingen, cat# 553708, 1:500), Rabbit anti-α-smooth-muscle-actin (Abcam, ab15734, 1:200), Rabbit anti-α-Actinin (Abcam, ab68167, 1:200), Mouse anti-α-Actinin(Sigma, A-7811, 1:500), Goat Anti-tdTomato (SICGEN, AB8181-200, 1:500). The following secondary antibodies were diluted 1:250 in 0.125%PBST with DAPI (Invitrogen, Catalog # D1306, 1:1000) and incubated for 1.5 hour at RT: Donkey Anti-Rabbit IgG-Alexa 488, 594, 647 (Invitrogen, Catalog # A32790, # A32754, # A32795); Donkey anti-Goat IgG-Alexa 594 (Invitrogen, Catalog # SA5-10088), Donkey anti-Mouse IgG-Alexa 488 (Invitrogen, Catalog # A32766), Goat anti-Chicken IgG-Alexa 488 (Invitrogen, Catalog # A32931), Donkey anti-Rat IgG-Alexa 488 (Invitrogen, Catalog # A-21208). All images were processed using Nikon NIS Elements software, ImageJ and Adobe Illustrator.

### Statistical Analysis of Clonal Events

To ensure that the clones we analyzed were the result of a single recombination event, we employed several different techniques including 1) using a low dose of tamoxifen, 2) utilizing the *Rosa26-Confetti* mouse line in which for each recombination event, only one of four different fluorophores is expressed, 3) separately analyzing unicolor embryos as well as multi-color embryos and 4) applying a rigorous statistical analysis based on the number of cells in a clone in order to exclude single color clones that may have resulted from two recombinant events. This statistical analysis involved embryos harvested at E9.5 in which single color clones were noted in both extraembryonic and cardiac regions. In these embryos, we calculated the number of cells in each clone. A mixture of two Gaussian distributions^64^ was fit to the data using expectation maximization. The distribution for the counts was then given by:

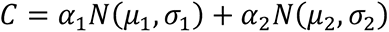

with:

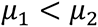

α_1_, α_2_ were the mixing parameters, and *N*(μ,σ) was a Gaussian distribution with mean μ and standard deviation σ. The first of the Gaussian distributions represented the cell count distribution from a single recombination event while the other represented the cell count distribution from multi-recombination events. This analysis was performed separately on E6.75 and E7.25 induced clones. Clones which had a likelihood of belonging to the Gaussian distribution with a smaller mean were consider clonal and the clones that fit the larger mean Gaussian distribution were considered multiclonal and were excluded from our analysis of multipotentiality.

### Statistics and reproducibility

Replicates and statistical tests are described in the figure legends. No statistical methods were used to predetermine sample size. Experiments did not employ randomization nor investigator blinding. All experimental results were analyzed with at least three independent embryos. Wilcoxon rank sum tests were used for statistical tests for differential gene expression analysis including the tests supporting the box and whisker plots. Markers were defined by Seurat’s default settings (at least >0.25 log fold increase over the opposing group, at most <0.01 unadjusted p-value, and at least >10% cells expressing). On the boxplots, a p-value < 0.01 was considered to be statistically significant as indicated by *. Box and whisker plots were created with standard parameters from ggplot2.

### Reporting summary

Further information on research design is available in the Nature Research Reporting Summary linked to this paper.

### Data and code availability

The scRNA-Seq data set supporting results of this article is available in the GEO database. Visualization of gene expression of the scRNA-seq is available on the UCSC cell browser at https://cells.ucsc.edu/. The R scripts are available upon request.

## Main References

1 Meilhac, S. M. & Buckingham, M. E. The deployment of cell lineages that form the mammalian heart. Nature reviews. Cardiology 15, 705–724, doi:10.1038/s41569-018-0086-9 (2018).

2 Meilhac, S. M., Esner, M., Kelly, R. G., Nicolas, J. F. & Buckingham, M. E. The clonal origin of myocardial cells in different regions of the embryonic mouse heart. Developmental cell 6, 685–698, doi:10.1016/s1534-5807(04)00133-9 (2004).

3 Devine, W. P., Wythe, J. D., George, M., Koshiba-Takeuchi, K. & Bruneau, B. G. Early patterning and specification of cardiac progenitors in gastrulating mesoderm. eLife 3, doi:10.7554/eLife.03848 (2014).

4 Lescroart, F. et al. Early lineage restriction in temporally distinct populations of Mesp1 progenitors during mammalian heart development. Nature cell biology 16, 829–840, doi:10.1038/ncb3024 (2014).

5 Cai, C. L. et al. Isl1 identifies a cardiac progenitor population that proliferates prior to differentiation and contributes a majority of cells to the heart. Developmental cell 5, 877–889, doi:10.1016/s1534-5807(03)00363-0 (2003).

6 Kelly, R. G., Brown, N. A. & Buckingham, M. E. The arterial pole of the mouse heart forms from Fgf10-expressing cells in pharyngeal mesoderm. Developmental cell 1, 435–440, doi:10.1016/s1534-5807(01)00040-5 (2001).

7 Prall, O. W. et al. An Nkx2-5/Bmp2/Smad1 negative feedback loop controls heart progenitor specification and proliferation. Cell 128, 947–959, doi:10.1016/j.cell.2007.01.042 (2007).

8 Bu, L. et al. Human ISL1 heart progenitors generate diverse multipotent cardiovascular cell lineages. Nature 460, 113–117, doi:10.1038/nature08191 (2009).

9 Lescroart, F. et al. Clonal analysis reveals common lineage relationships between head muscles and second heart field derivatives in the mouse embryo. Development (Cambridge, England) 137, 3269–3279, doi:10.1242/dev.050674 (2010).

10 Lescroart, F. et al. Clonal analysis reveals a common origin between nonsomite-derived neck muscles and heart myocardium. Proceedings of the National Academy of Sciences of the United States of America 112, 1446–1451, doi:10.1073/pnas.1424538112 (2015).

11 Diogo, R. et al. A new heart for a new head in vertebrate cardiopharyngeal evolution. Nature 520, 466–473, doi:10.1038/nature14435 (2015).

12 Liang, X. et al. HCN4 dynamically marks the first heart field and conduction system precursors. Circulation research 113, 399–407, doi:10.1161/circresaha.113.301588 (2013).

13 Später, D. et al. A HCN4+ cardiomyogenic progenitor derived from the first heart field and human pluripotent stem cells. Nature cell biology 15, 1098–1106, doi:10.1038/ncb2824 (2013).

14 Komiyama, M., Ito, K. & Shimada, Y. Origin and development of the epicardium in the mouse embryo. Anatomy and embryology 176, 183–189, doi:10.1007/bf00310051 (1987).

15 Mikawa, T. & Gourdie, R. G. Pericardial mesoderm generates a population of coronary smooth muscle cells migrating into the heart along with ingrowth of the epicardial organ. Developmental biology 174, 221–232, doi:10.1006/dbio.1996.0068 (1996).

16 Gittenberger-de Groot, A. C., Vrancken Peeters, M. P., Mentink, M. M., Gourdie, R. G. & Poelmann, R. E. Epicardium-derived cells contribute a novel population to the myocardial wall and the atrioventricular cushions. Circulation research 82, 1043–1052, doi:10.1161/01.res.82.10.1043 (1998).

17 Grieskamp, T., Rudat, C., Lüdtke, T. H., Norden, J. & Kispert, A. Notch signaling regulates smooth muscle differentiation of epicardium-derived cells. Circulation research 108, 813–823, doi:10.1161/circresaha.110.228809 (2011).

18 Maya-Ramos, L., Cleland, J., Bressan, M. & Mikawa, T. Induction of the Proepicardium. Journal of developmental biology 1, 82–91, doi:10.3390/jdb1020082 (2013).

19 Saga, Y. et al. MesP1 is expressed in the heart precursor cells and required for the formation of a single heart tube. Development (Cambridge, England) 126, 3437–3447 (1999).

20 Madisen, L. et al. A robust and high-throughput Cre reporting and characterization system for the whole mouse brain. Nature neuroscience 13, 133–140, doi:10.1038/nn.2467 (2010).

21 Oginuma, M., Hirata, T. & Saga, Y. Identification of presomitic mesoderm (PSM)-specific Mesp1 enhancer and generation of a PSM-specific Mesp1/Mesp2-null mouse using BAC-based rescue technology. Mechanisms of development 125, 432–440, doi:10.1016/j.mod.2008.01.010 (2008).

22 Yoshida, T., Vivatbutsiri, P., Morriss-Kay, G., Saga, Y. & Iseki, S. Cell lineage in mammalian craniofacial mesenchyme. Mechanisms of development 125, 797–808, doi:10.1016/j.mod.2008.06.007 (2008).

23 Harel, I. et al. Distinct origins and genetic programs of head muscle satellite cells. Developmental cell 16, 822–832, doi:10.1016/j.devcel.2009.05.007 (2009).

24 Chan, S. S. et al. Mesp1 patterns mesoderm into cardiac, hematopoietic, or skeletal myogenic progenitors in a context-dependent manner. Cell stem cell 12, 587–601, doi:10.1016/j.stem.2013.03.004 (2013).

25 Butler, A., Hoffman, P., Smibert, P., Papalexi, E. & Satija, R. Integrating single-cell transcriptomic data across different conditions, technologies, and species. Nature biotechnology 36, 411–420, doi:10.1038/nbt.4096 (2018).

26 Scialdone, A. et al. Resolving early mesoderm diversification through single-cell expression profiling. Nature 535, 289–293, doi:10.1038/nature18633 (2016).

27 Pijuan-Sala, B. et al. A single-cell molecular map of mouse gastrulation and early organogenesis. Nature 566, 490–495, doi:10.1038/s41586-019-0933-9 (2019).

28 Farrell, J. A. et al. Single-cell reconstruction of developmental trajectories during zebrafish embryogenesis. Science (New York, N.Y.) 360, doi:10.1126/science.aar3131 (2018).

29 Ueno, H. & Weissman, I. L. Clonal analysis of mouse development reveals a polyclonal origin for yolk sac blood islands. Developmental cell 11, 519–533, doi:10.1016/j.devcel.2006.08.001 (2006).

30 Tam, P. P. The allocation of cells in the presomitic mesoderm during somite segmentation in the mouse embryo. Development (Cambridge, England) 103, 379–390 (1988).

31 Gouti, M. et al. A Gene Regulatory Network Balances Neural and Mesoderm Specification during Vertebrate Trunk Development. Developmental cell 41, 243–261.e247, doi:10.1016/j.devcel.2017.04.002 (2017).

32 Harel, I. et al. Pharyngeal mesoderm regulatory network controls cardiac and head muscle morphogenesis. Proceedings of the National Academy of Sciences of the United States of America 109, 18839–18844, doi:10.1073/pnas.1208690109 (2012).

33 Bao, Z. Z., Bruneau, B. G., Seidman, J. G., Seidman, C. E. & Cepko, C. L. Regulation of chamber-specific gene expression in the developing heart by Irx4. Science (New York, N.Y.) 283, 1161–1164, doi:10.1126/science.283.5405.1161 (1999).

34 Bruneau, B. G. et al. Chamber-specific cardiac expression of Tbx5 and heart defects in Holt-Oram syndrome. Developmental biology 211, 100–108, doi:10.1006/dbio.1999.9298 (1999).

35 van den Berg, G. et al. A caudal proliferating growth center contributes to both poles of the forming heart tube. Circulation research 104, 179–188, doi:10.1161/circresaha.108.185843 (2009).

36 Barnes, R. M. et al. MEF2C regulates outflow tract alignment and transcriptional control of Tdgf1. Development (Cambridge, England) 143, 774–779, doi:10.1242/dev.126383 (2016).

37 Rudat, C. et al. Upk3b is dispensable for development and integrity of urothelium and mesothelium. PloS one 9, e112112, doi:10.1371/journal.pone.0112112 (2014).

38 Facucho-Oliveira, J., Bento, M. & Belo, J. A. Ccbe1 expression marks the cardiac and lymphatic progenitor lineages during early stages of mouse development. The International journal of developmental biology 55, 1007–1014, doi:10.1387/ijdb.113394jf (2011).

39 Fujii, M. et al. Sfrp5 identifies murine cardiac progenitors for all myocardial structures except for the right ventricle. Nature communications 8, 14664, doi:10.1038/ncomms14664 (2017).

40 Saito, Y., Kojima, T. & Takahashi, N. Mab21l2 is essential for embryonic heart and liver development. PloS one 7, e32991, doi:10.1371/journal.pone.0032991 (2012).

41 Kraus, F., Haenig, B. & Kispert, A. Cloning and expression analysis of the mouse T-box gene Tbx18. Mechanisms of development 100, 83–86, doi:10.1016/s0925-4773(00)00494-9 (2001).

42 de Soysa, T. Y. et al. Single-cell analysis of cardiogenesis reveals basis for organ-level developmental defects. Nature 572, 120–124, doi:10.1038/s41586-019-1414-x (2019).

43 Lupu, I. E., Redpath, A. N. & Smart, N. Spatiotemporal Analysis Reveals Overlap of Key Proepicardial Markers in the Developing Murine Heart. Stem cell reports 14, 770–787, doi:10.1016/j.stemcr.2020.04.002 (2020).

44 Di Bella, D. J. et al. Molecular Logic of Cellular Diversification in the Mammalian Cerebral Cortex. bioRxiv, 2020.2007.2002.185439, doi:10.1101/2020.07.02.185439 (2020).

45 Riley, P., Anson-Cartwright, L. & Cross, J. C. The Hand1 bHLH transcription factor is essential for placentation and cardiac morphogenesis. Nature genetics 18, 271–275, doi:10.1038/ng0398-271 (1998).

46 Firulli, A. B., McFadden, D. G., Lin, Q., Srivastava, D. & Olson, E. N. Heart and extra-embryonic mesodermal defects in mouse embryos lacking the bHLH transcription factor Hand1. Nature genetics 18, 266–270, doi:10.1038/ng0398-266 (1998).

47 Srivastava, D., Cserjesi, P. & Olson, E. N. A subclass of bHLH proteins required for cardiac morphogenesis. Science (New York, N.Y.) 270, 1995–1999, doi:10.1126/science.270.5244.1995 (1995).

48 Merscher, S. et al. TBX1 is responsible for cardiovascular defects in velo-cardio-facial/DiGeorge syndrome. Cell 104, 619–629, doi:10.1016/s0092-8674(01)00247-1 (2001).

49 Zhang, Z., Huynh, T. & Baldini, A. Mesodermal expression of Tbx1 is necessary and sufficient for pharyngeal arch and cardiac outflow tract development. Development (Cambridge, England) 133, 3587–3595, doi:10.1242/dev.02539 (2006).

50 Lin, Q., Schwarz, J., Bucana, C. & Olson, E. N. Control of mouse cardiac morphogenesis and myogenesis by transcription factor MEF2C. Science (New York, N.Y.) 276, 1404–1407, doi:10.1126/science.276.5317.1404 (1997).

51 Jongbloed, M. R. et al. Expression of Id2 in the second heart field and cardiac defects in Id2 knock-out mice. Developmental dynamics : an official publication of the American Association of Anatomists 240, 2561–2577, doi:10.1002/dvdy.22762 (2011).

52 Moskowitz, I. P. et al. A molecular pathway including Id2, Tbx5, and Nkx2-5 required for cardiac conduction system development. Cell 129, 1365–1376, doi:10.1016/j.cell.2007.04.036 (2007).

53 Fraidenraich, D. et al. Rescue of cardiac defects in id knockout embryos by injection of embryonic stem cells. Science (New York, N.Y.) 306, 247–252, doi:10.1126/science.1102612 (2004).

54 Bamforth, S. D. et al. Cited2 controls left-right patterning and heart development through a Nodal-Pitx2c pathway. Nature genetics 36, 1189–1196, doi:10.1038/ng1446 (2004).

55 Barnes, R. M. et al. Hand2 loss-of-function in Hand1-expressing cells reveals distinct roles in epicardial and coronary vessel development. Circulation research 108, 940–949, doi:10.1161/circresaha.110.233171 (2011).

56 Biben, C. & Harvey, R. P. Homeodomain factor Nkx2-5 controls left/right asymmetric expression of bHLH gene eHand during murine heart development. Genes & development 11, 1357–1369, doi:10.1101/gad.11.11.1357 (1997).

57 Thomas, T., Yamagishi, H., Overbeek, P. A., Olson, E. N. & Srivastava, D. The bHLH factors, dHAND and eHAND, specify pulmonary and systemic cardiac ventricles independent of left-right sidedness. Developmental biology 196, 228–236, doi:10.1006/dbio.1998.8849 (1998).

58 Robinson, S. P., Langan-Fahey, S. M., Johnson, D. A. & Jordan, V. C. Metabolites, pharmacodynamics, and pharmacokinetics of tamoxifen in rats and mice compared to the breast cancer patient. Drug metabolism and disposition: the biological fate of chemicals 19, 36–43 (1991).

59 Wilson, C. H. et al. The kinetics of ER fusion protein activation in vivo. Oncogene 33, 4877–4880, doi:10.1038/onc.2014.78 (2014).

60 Aanhaanen, W. T. et al. The Tbx2+ primary myocardium of the atrioventricular canal forms the atrioventricular node and the base of the left ventricle. Circulation research 104, 1267–1274, doi:10.1161/circresaha.108.192450 (2009).

61 Li, G. et al. Single cell expression analysis reveals anatomical and cell cycle-dependent transcriptional shifts during heart development. Development (Cambridge, England) 146, doi:10.1242/dev.173476 (2019).

62 Harrelson, Z. et al. Tbx2 is essential for patterning the atrioventricular canal and for morphogenesis of the outflow tract during heart development. Development (Cambridge, England) 131, 5041–5052, doi:10.1242/dev.01378 (2004).

63 Snippert, H. J. et al. Intestinal crypt homeostasis results from neutral competition between symmetrically dividing Lgr5 stem cells. Cell 143, 134–144, doi:10.1016/j.cell.2010.09.016 (2010).

64 Benaglia, T., Chauveau, D., Hunter, D., R. & Young, D., S. mixtools: An R Package for Analyzing Finite Mixture Models. Journal of Statistical Software 32, 1–29 (2009).

65 Nowotschin, S. et al. The emergent landscape of the mouse gut endoderm at single-cell resolution. Nature 569, 361–367, doi:10.1038/s41586-019-1127-1 (2019).

66 Moretti, A. et al. Multipotent embryonic isl1+ progenitor cells lead to cardiac, smooth muscle, and endothelial cell diversification. Cell 127, 1151–1165, doi:10.1016/j.cell.2006.10.029 (2006).

67 Sun, Y. et al. Islet 1 is expressed in distinct cardiovascular lineages, including pacemaker and coronary vascular cells. Developmental biology 304, 286–296, doi:10.1016/j.ydbio.2006.12.048 (2007).

68 Orkin, S. H. & Zon, L. I. Hematopoiesis: an evolving paradigm for stem cell biology. Cell 132, 631–644, doi:10.1016/j.cell.2008.01.025 (2008).

69 Kwon, G. S., Viotti, M. & Hadjantonakis, A. K. The endoderm of the mouse embryo arises by dynamic widespread intercalation of embryonic and extraembryonic lineages. Developmental cell 15, 509–520, doi:10.1016/j.devcel.2008.07.017 (2008).

70 Kruithof, B. P. et al. BMP and FGF regulate the differentiation of multipotential pericardial mesoderm into the myocardial or epicardial lineage. Developmental biology 295, 507–522, doi:10.1016/j.ydbio.2006.03.033 (2006).

71 Van Handel, B. et al. Scl represses cardiomyogenesis in prospective hemogenic endothelium and endocardium. Cell 150, 590–605, doi:10.1016/j.cell.2012.06.026 (2012).

72 Zhou, B. et al. Adult mouse epicardium modulates myocardial injury by secreting paracrine factors. The Journal of clinical investigation 121, 1894–1904, doi:10.1172/jci45529 (2011).

73 Smart, N. et al. De novo cardiomyocytes from within the activated adult heart after injury. Nature 474, 640–644, doi:10.1038/nature10188 (2011).

74 Zangi, L. et al. Insulin-Like Growth Factor 1 Receptor-Dependent Pathway Drives Epicardial Adipose Tissue Formation After Myocardial Injury. Circulation 135, 59–72, doi:10.1161/circulationaha.116.022064 (2017).

75 Matthiesen, N. B. et al. Congenital Heart Defects and Indices of Placental and Fetal Growth in a Nationwide Study of 924 422 Liveborn Infants. Circulation 134, 1546–1556, doi:10.1161/circulationaha.116.021793 (2016).

## Method References

64 Benaglia, T., Chauveau, D., Hunter, D. R. & Young, D., S. mixtools: An R Package for Analyzing Finite Mixture Models. Journal of Statistical Software 32, 1–29 (2009).

76 Yao, X. et al. Tild-CRISPR Allows for Efficient and Precise Gene Knockin in Mouse and Human Cells. Developmental cell 45, 526–536.e525, doi:10.1016/j.devcel.2018.04.021 (2018).

77 Downs, K. M. & Davies, T. Staging of gastrulating mouse embryos by morphological landmarks in the dissecting microscope. Development (Cambridge, England) 118, 1255–1266 (1993).

78 Harris, M. A. et al. The Gene Ontology (GO) database and informatics resource. Nucleic acids research 32, D258–261, doi:10.1093/nar/gkh036 (2004).

